# RPI-Zero: Zero-shot RNA-Protein Interaction Prediction

**DOI:** 10.64898/2026.11.04.764286

**Authors:** Kerim Eren, E. İrem Esendemir, Sobhan Shukueian Tabrizi, Helyasadat Hashemi Aghdam, A. Ercüment Çiçek

## Abstract

In vivo protein–RNA interactions are mapped almost exclusively by CLIP-seq, which profiles one protein in one cell type per experiment and needs a specific antibody; only about 10% of the roughly 2,000 human RNA-binding proteins have been assayed, and the Protein Data Bank holds around 5,000 protein–RNA complexes, a quarter of them ribosomes. Sequence-based prediction could extend coverage to unprofiled, newly discovered, or designed molecules, but existing predictors are not built for that case. One family trains a separate model per protein and cannot move beyond proteins with CLIP data; the other scores pairs but targets a single RNA class. Most are evaluated with random splits and randomly paired negatives that leave near-identical sequences on both sides of the split, and many require structure or read only short windows, discarding the context around a binding site. Their accuracy falls sharply when whole molecules are held out.

We address this with two contributions. First, a strict benchmark of 2.1 million examples built from 11.6 million peaks of CLIP experiments targeting 209 proteins, taken from four public databases, in which each RNA is a spliced transcript with nucleotide-level labels, and sequence-similarity clusters are assigned whole to training, validation, and test sets, giving separate slices for unseen RNAs, unseen proteins, and unseen pairs. Negatives are matched to positives in length and in the number of other proteins binding the transcript, removing the shortcut by which a model predicts binding from how popular a transcript is without reading the protein. Second, RPI-Zero, a sequence-only model that reads any full-length protein and RNA jointly in up to 4,096 tokens, so the RNA representation is conditioned on the protein, and describes every protein as a mixture of shared motif prototypes, placing unseen proteins in the same terms as training ones. One model covers all proteins and RNA classes, uses no structure, and needs no interaction data for a new molecule. Compared with five state-of-the-art methods, it achieves the highest score in eight different metrics on unseen RNAs, unseen proteins, and unseen pairs. The model and data are available at https://github.com/ciceklab/RPI-Zero and https://zenodo.org/records/23748141, respectively.

## 1 Introduction

In vivo protein–RNA interactions are detected primarily by crosslinking and immunoprecipitation followed by sequencing techniques (CLIP-seq), which covalently trap protein–RNA contacts with ultraviolet light, isolate the protein of interest with an antibody, and sequence the bound RNA fragments [11, 15, 18, 37, 39]. Although CLIP-seq captures binding in cellular context, each experiment involves a single protein in a single cell type, requires a highly specific antibody that is unavailable for many RNA-binding proteins, and is costly to perform [10, 16, 35, 38]. Therefore, only a small fraction (10%) of the roughly 2,000 human RNA-binding proteins have been profiled in a few cell lines [48]. Structural approaches face the opposite constraint, resolving interfaces at atomic detail but only for complexes stable enough to crystallize or image, which excludes most flexible RNA interactions. The Protein Data Bank [4] contains only about 5,000 protein–RNA complex structures, and a quarter of them are ribosomes [3]. This motivates computational methods that can infer protein–RNA interfaces directly from sequence, at scale and can generalize to unseen targets such as synthetic proteins or RNA molecules, without the need for structures or targeted experiments.

There is a vast literature on methods that predict RNA-protein interactions which can be roughly classified into two families. The first set of models train a predictor model per protein and optionally per cell line [9, 23, 34, 52] to predict if a given RNA binds to it. However, this means that they are specific for proteins for which CLIP-seq data exist and cannot be generalized beyond. The second set of methods build a single model that takes a pair of RNA and protein molecules as input and predicts whether they bind. However, most of such methods are specific to a class of RNA such as lncRNAs [17, 22, 43, 51], other non-coding RNAs (ncRNAs) [12, 28, 31, 40, 46], circRNAs [41], microRNAs [2], and mRNAs [48].

The reported accuracy of many of these methods does not carry over to unseen data. Most were tested with k-fold cross-validation on small data sets, often built from PDB complexes [26–28, 31, 40, 46, 47]. Their negative pairs were mostly made by random pairing [12, 17, 26–28, 31, 40, 43, 47]. Such splits put similar sequences in both the training set and the test set, so a model can score well by memorizing [5]. Graph and matrix methods also depend on the known interactions around a pair. NPI-GNN reaches almost the same accuracy without sequence information, and its authors state that it relies heavily on the network around a pair [31]. The accuracy of ncRPI-LGAT drops when this network is sparse [12]. The authors of MFNN, which uses only an interaction matrix, state that it cannot score a new circRNA or RBP [41]. Studies that hold out whole molecules report lower accuracy. In LPI-deepGBDT, performance was much higher when random pairs were held out than when whole lncRNAs or whole proteins were held out, and it was lowest for held-out proteins [51]. When no test RNA or protein was similar to the training set, the performance of four state-of-the-art methods evaluated in a recent benchmark study degraded substantially, and two of them predicted only one class [5]. Yu et al. [48] held out whole proteins. The accuracy of their own model dropped, and most published tools were only slightly better than random. It is therefore unclear how well most methods work on a new protein, a new RNA, or a new pair. Our model is built for this case. We test it on sets in which the RNA, the protein, or both are dissimilar to the training data.

Many methods also need structure as input. Several use the secondary structure of the RNA predicted from its sequence [17, 22, 23, 28, 40, 43, 47]. Some add features of the protein structure, such as secondary structure, contact maps, or 3D structure [2, 17, 22, 36, 40, 43]. Experimental structures are not available for most of the RNAs and proteins [20, 36], and predicted RNA structures often differ from the structures experimentally obtained in cells [34]. For example, in PrismNet, adding a predicted structure to the sequence gave about the same performance as the sequence alone, and only the actual structure improved the performance [34]. However, an in silico designed RNA would not have a determined structure, and even if it did, the flexibility of RNA hinders the usefulness of this information, as it can attain various different conformations with similar energies. The predicted structures of proteins also might not help this task. Yu et al. [48] used AlphaFold-predicted structure features of the target protein, and this did not close the performance gap for unseen proteins. To enable our model to work in zero-shot settings, we only use the sequence of the RNA and the protein.

The input length is another limit. Methods that predict binding sites read short windows. DeepCLIP was trained on sites of 12 to 75 nt, and PrismNet, HDRNet, and DeepMiRBP use RNA windows of 101 nt [2, 9, 34, 52]. The HDRNet authors note that this length restriction makes precise prediction difficult [52]. Pair-level methods that use large pretrained models face limits on the input as well. ZHMolGraph is limited by the 1,022 nt input of RNA-FM [6], so it cuts long RNAs into segments and averages the segment embeddings [20]. CORAL removes proteins longer than 1,250 residues and RNAs longer than 6,000 nt because of GPU memory limitation [5]. Structure prediction adds a limit of its own. lncPro uses RNAsubopt, which processes only the first 4,095 nt of an RNA [22, 27, 46]. EDLMFC keeps only RNAs of up to 500 nt because SPOT-RNA works only up to that length [40]. K-mer methods accept any length, but k-mer counts do not record where on the sequence a k-mer lies [17, 26–28, 40, 43, 47, 51]. In our data, the median CLIP peak is 35 nt long and the median transcript is 2,023 nt long, so a short window sees only a small part of the context of a binding site. Our model reads the full-length protein and a transcript in a single input. The transcript can be up to 3,966 nt long for the shortest protein in our data and up to 1,289 nt for the longest. A longer transcript is cut into overlapping chunks.

We make two contributions to address these problems. First, we introduce a benchmark for RNA-protein interaction prediction that measures how well a model works on new molecules. Any model that takes an RNA sequence and a protein sequence can be trained and tested on it. We built the benchmark from 11.6 million CLIP peaks that target 209 proteins, taken from four public databases [38, 44, 49, 53]. Each RNA in the benchmark is a spliced canonical transcript instead of a short peak sequence. We cluster the transcripts and the proteins by sequence similarity [33] and place whole clusters in the training, validation, or test sets. The validation set and the test set therefore have separate slices for unseen RNAs, unseen proteins, and unseen pairs. We match the negative examples to the positives in terms of length and in the number of other proteins that bind the transcript so that models cannot learn shortcuts. The benchmark contains 2.1 million examples (see Section 2.1).

Second, we propose RPI-Zero, a model that predicts the binding probability of an RNA and a protein using only their sequences, enabling it to work with any RNA and protein of interest. RPI-Zero uses the embeddings of an RNA-interaction foundation model [32], which returns a joint embedding for the protein and the RNA of up to 4,096 tokens. It assigns each nucleotide a binding score, and a pooling of these scores gives the binding probability of the pair. The model uses the full canonical transcripts with CLIP-peak labels in binding regions, which enable the model to understand the binding region within the context of the full sequence. We compare RPI-Zero with five state-of-the-art methods that take the sequences of an RNA and a protein as input and can read full-length transcripts [5, 7, 20, 27, 31]. We retrain all of them on our benchmark and test every method on the same three slices. RPI-Zero has the highest AUPRC on all three slices, with 0.753 on unseen RNAs, 0.637 on unseen proteins, and 0.589 on unseen pairs. This is 2.7%, 7.4%, and 4.6% higher than the AUPRC of the second-best method, which is a different method on each slice (ZHMolTopoRPI, IPMiner, and CORAL). RPI-Zero also has the highest AUROC, accuracy, F1, and MCC on every slice. The model and the data are available at https://github.com/ciceklab/RPI-Zero and https://zenodo.org/records/23748141, respectively.

## 2 Methods

### 2.1 Dataset

We design each step so that a model scores well only if it learns how RNAs and proteins bind, and not from how the data was built (Fig. 1, Supplementary Section 1).

**Fig. 1.**
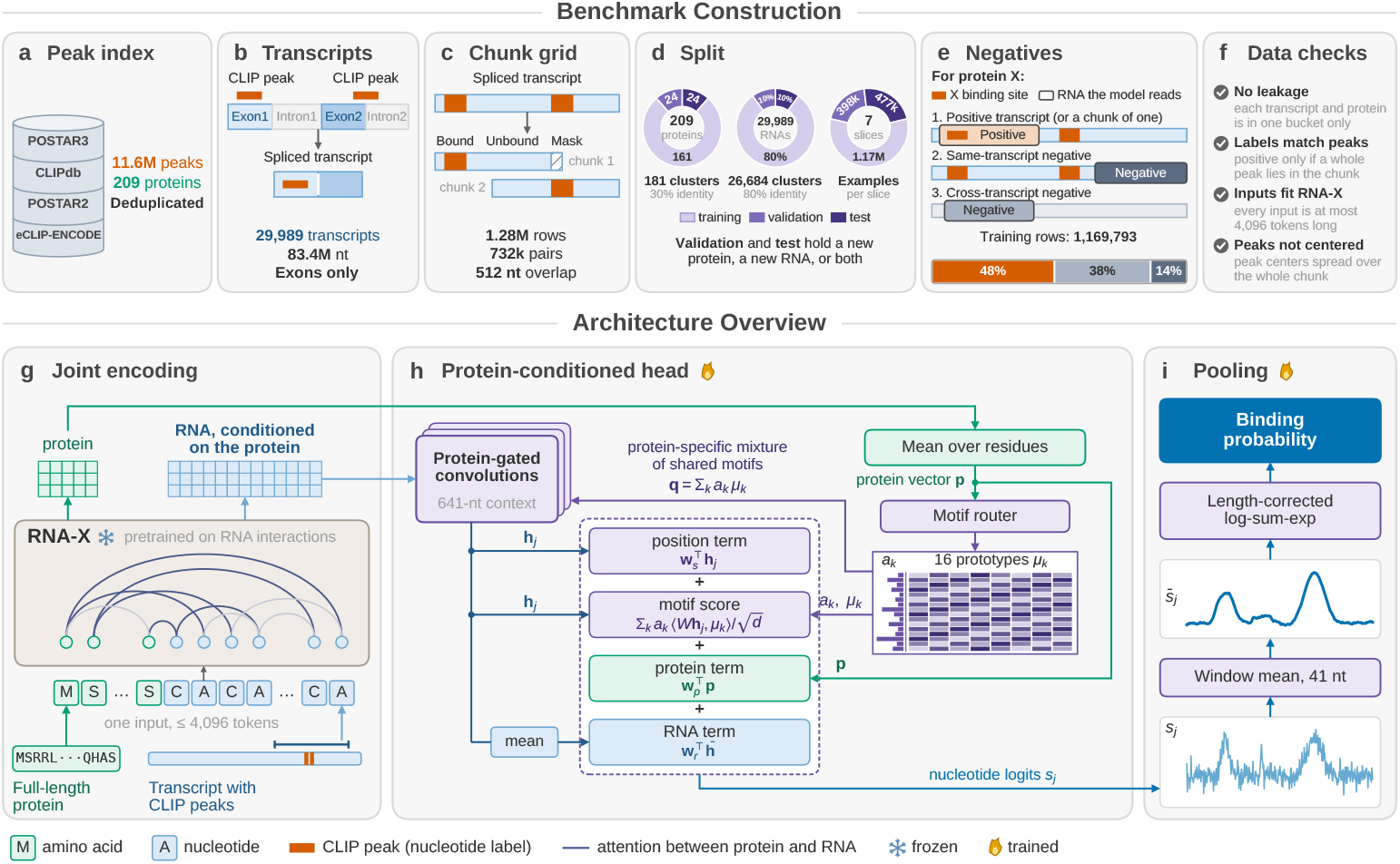
Overview of the benchmark (a–f) and of RPI-Zero (g–i). (a) We merge the data of CLIP experiments from four databases. This gives 11.6 million peaks bound by 209 proteins. (b) For each peak, we take the canonical transcript from the genome and use its spliced sequence, which holds only the exons. We keep a peak only if it lies fully inside one exon. (c) If a transcript is too long to fit into the input of RNA-X together with the protein, we cut it into chunks that overlap by 512 nt. The chunks lie on a fixed grid that does not depend on where the peaks are. Nucleotides inside a peak are labeled bound, and the others unbound. If a chunk border cuts a peak, we mask the cut piece and leave it out of the loss. (d) Whole clusters of proteins and transcripts are assigned to buckets, which give seven slices (Table 1). (e) The three kinds of examples for a protein X, and their shares of the training slice. The RNA of an example is a transcript or a chunk of one. A positive example holds a whole peak bound by X. A same-transcript negative is a part of a transcript that X binds, with no whole peak bound by X. A cross-transcript negative comes from a transcript that X does not bind. (f) Four checks that we run on all examples (Supplementary Section 1.6). The last one shows that the position of a peak does not reveal the label. (g) Frozen RNA-X reads the full protein and the RNA as one input. (h) The trained head describes the protein as a mixture of 16 shared motif prototypes. The mixture gates convolutions over the RNA, and the logit of each nucleotide is the sum of four terms (Section 2.2). (i) A window mean over 41 nt and a length-corrected log-sum-exp give the binding probability. The tracks are illustrations.

#### 2.1.1 Transcripts and labels

We take the CLIP peaks of four databases, CLIPdb [44], ENCODE eCLIP [38], POSTAR3 [49], and POSTAR2 [53]. Each CLIP experiment targets one human RNA-binding protein, and a peak is an RNA region that this protein binds, given as an interval on the hg38 genome. The merged set has 11.6 million peaks bound by 209 proteins (Fig. 1a), and we represent each protein by its amino acid sequence.

**Table 1.** Details of the benchmark slices. Here, a bucket for a molecule can be training, validation or test. All molecules in a sequence cluster in the benchmark are placed as a whole in a single bucket (never separated; Section 2.1). The RNA bucket and the protein bucket of an interaction instance (example) determine its slice (unseen RNA, protein or pair). In an unseen RNA/protein slice, the RNA/protein of each example is new to the model, respectively. In an unseen pair slice, both are new. In a validation slice, the unseen molecules come from the validation bucket and the seen molecules from the training bucket. In a test slice, the unseen molecules come from the test bucket and the seen molecules from the training or validation bucket. Models are trained on the training slice and selected on the validation slices, so validation molecules count as seen in the test slices. Each cell gives the slice and, below it, its number of examples / positives. Examples: protein–RNA pairs, where the RNA is a transcript or a chunk of one. Positives: examples whose RNA holds at least one whole CLIP peak bound by the protein. A slice that covers two cells is written in both, and its numbers count the whole slice once.

| **Protein**<br>**RNA** | **Training** | **Validation** | **Test** |
|---|---|---|---|
| **Training** | Training<br>1,169,793 / 562,637 | Validation (unseen protein)<br>221,288 / 96,668 | Test (unseen protein)<br>269,408 / 117,336 |
| **Validation** | Validation (unseen RNA)<br>149,016 / 71,906 | Validation (unseen pair)<br>27,307 / 11,903 | Test (unseen protein)<br>269,408 / 117,336 |
| **Test** | Test (unseen RNA)<br>177,838 / 83,584 | Test (unseen RNA)<br>177,838 / 83,584 | Test (unseen pair)<br>29,808 / 12,830 |

We use the full spliced transcript as the RNA, so that the model sees the context of a binding site. For each gene with a peak, we take one transcript of GENCODE release 50 [25], preferring the MANE Select transcript [24], then the Ensembl canonical transcript [45], and then the longest one. This gives 29,989 transcripts, and all proteins see the same transcript of a gene. We keep a peak only if it lies fully inside one exon of the chosen transcript (Fig. 1b). Because the input and the labels are both on the spliced transcript, the model can be fine-tuned further on other RNA–protein databases that give the RNA as a transcript, such as NPInter v5.0 [50]. Nucleotides inside a peak have the label 1, and all others have the label 0. Binding in introns and across exon junctions is therefore not part of the benchmark.

We use RNA-X [32] as the RNA interaction foundation model to supply the embeddings. This model reads at most 4,096 tokens, which the protein and the RNA share, so the RNA can have at most 4,093 ™ *L*_*p*_ nt for a protein of *L*_*p*_ amino acids. We cut a transcript longer than this into parts, which we call chunks in the rest of the paper. The chunks overlap by 512 nt and lie on a fixed grid from the 5^′^ end (Fig. 1c). The grid does not depend on the peaks, because a chunk centered on a peak would reveal the label through the position of the peak. If a chunk border cuts a peak, we mask the cut piece, because the chunk does not show the whole site. An example is a protein paired with a transcript or a chunk of one, and it is positive if its RNA holds at least one whole peak bound by the protein.

#### 2.1.2 Clustering and data splits

We cluster whole transcripts and not peak sequences, because the model reads whole transcripts, and two peaks on the same transcript can fall into different clusters of peak sequences. We cluster the transcripts at 80% and the proteins at 30% sequence identity with MMseqs2 [33], and we put each cluster in a single bucket (Fig. 1d). The training, validation, and test buckets hold 80%, 10%, and 10% of the examples for the transcripts, and of the peaks for the proteins. We do not train on 637 transcripts of the training bucket, and we use their examples with training proteins only to follow training (Supplementary Section 1.4). Table 1 gives the slices.

#### 2.1.3 Negative examples

We generate two kinds of negative examples (Fig. 1e). Same-transcript negatives come from transcripts to which the protein binds, but only parts (chunks) of these transcripts that do not have a CLIP peak are used. Because they originate from the same transcripts as the positives, the gene, its expression level, and its sequence composition highly resemble the positives and these cannot create shortcuts for the model.

Cross-transcript negatives originate from transcripts that the protein does not bind to. We call a transcript unbound (negative) only if no peak bound by the protein lies anywhere on its genomic span, including introns. This ensures that a negative transcript cannot hold a positive site in another example. We draw each negative from the same RNA bucket as the positives, so a negative is always in the same slice as its positives. Cross-transcript negatives can create a shortcut. A transcript that many proteins bind is rarely unbound, so unbound transcripts drawn at random have fewer binding partners than the positives. A model could then predict interactions from the *popularity* of a transcript regardless of the partner. We therefore jointly match (1) the popularity and (2) the length of the negative transcripts to that of positive partners for each protein. The benchmark contains 2,074,753 examples, of which 971,450 are positive.

### 2.2 Architecture

RPI-Zero uses RNA-X to compute an embedding for the RNA-protein pair [32]. It is a transformer with 14 layers and 136 million parameters, pretrained on over 100 million RNA interactions with proteins, RNA, and DNA. It reads the full protein and the RNA in one input (Fig. 1g), so the embedding of each nucleotide already depends on the protein. We keep RNA-X frozen and split its output into the protein embeddings 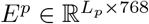 and the embeddings 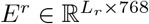 of an RNA of length *L*_*r*_.

The head maps the mean of the rows of *E*^*p*^ to a protein vector **p** ∈ ℝ^*d*^, with *d* = 256 (Fig. 1h). It holds *K* = 16 learned motif prototypes *µ*_*k*_ ∈ ℝ^*d*^ that all proteins share, and a router weights them for each protein with **a** = softmax(*W*_route_ **p**), which gives the protein context **q** = Σ_*k*_ *a*_*k*_ *µ*_*k*_. An unseen protein is described by the same 16 vectors as the training proteins, and only its weights are new.

The head maps each row of *E*^*r*^ to width *d* and passes it through five residual blocks. Each block applies a depthwise convolution of width 129 nt and multiplies its output by a sigmoid gate that depends on the nucleotide and on the protein context **q** [8]. The same RNA therefore gives different features for different proteins. After five blocks, the representation **h**_*j*_ of nucleotide *j* depends on a window of 641 nt around it. The logit of nucleotide *j* is 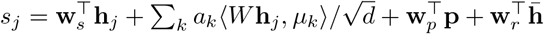, where *W* is a learned matrix and 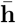 is the mean of **h**_*j*_. The motif score (second term) is high when the nucleotide matches the prototypes that the protein selects. The last two terms are constant within an example.

The pooling averages the logits over a centered window of 41 nt, which gives 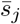, and then applies a log-sum-exp with a learned temperature *τ* and a length correction, 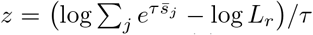 (Fig. 1i). The pair logit is *g* = *w z* + *b* with two learned scalars, and the binding probability is *σ*(*g*). The log-sum-exp ignores the order of the nucleotides, so the window mean is needed to favor runs of high logits, such as a CLIP peak, over scattered ones. We set the window to 41 nt, close to the length of a CLIP peak, which is 35 nt at the median and 47 nt on average in the training slice. The term ™log *L*_*r*_ keeps the score from growing with the RNA length, which depends on the protein through 4,093 ™ *L*_*p*_. Supplementary Section 2 gives all equations and hyperparameters.

### 2.3 Training

We train the motif-prototype head, which has 2 million parameters, with the loss *ℒ* = *ℒ*_nt_ + *λ*(*t*) *ℒ*_pair_, where *t* is the optimizer step. *ℒ*_nt_ is the binary cross-entropy of the nucleotide logits *s*_*j*_ against the peak labels, without padding and the masked pieces of cut peaks. Only 3.9% of the labeled training nucleotides lie in a peak, so we weight positive nucleotides by 49.06, which we selected by a hyperparameter search (Supplementary Section 2.9). *ℒ*_pair_ is the binary cross-entropy of the pair logit *g. ℒ*_nt_ gives the head a dense training signal at the positions of the peaks, which the pair label alone does not give. We train with *ℒ*_nt_ alone for the first 15,400 steps, so that the nucleotide logits learn where the peaks are, and then *λ* grows linearly to 10 over the next 15,400 steps (Supplementary Section 2). In an ablation study, removing the convolution blocks, *ℒ*_nt_, or *ℒ*_pair_ lowers the protein-macro AUROC on all three validation slices (Supplementary Section 4.2).

We balance the proteins and the labels in each batch. The number *n*_*p*_ of training examples per protein ranges from 99 to 32,238. We give protein *p* a number of draws proportional to 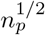, so that a few proteins do not dominate the gradient and rare proteins are not repeated too often. Half of the draws of each protein are positives, so that the pair logit cannot learn how often a protein binds instead of whether the RNA binds it. We draw without replacement for 154,000 steps, so every training example is drawn at least once.

### 2.4 Experimental setup

Each method is trained on the training slice and selected on the validation slices, which we also use to choose the hyperparameters of RPI-Zero. The test slices are scored once, after the checkpoint and the decision threshold of a method are fixed. Training RPI-Zero takes 12 hours on one NVIDIA RTX PRO 6000 GPU. We used protein-macro AUROC for the validation metric. For each protein with at least 10 positive and 10 negative examples in a slice, we compute the AUROC of a method on the examples of this protein. We then take the unweighted mean over these proteins, so every protein counts the same. The protein-macro AUPRC is computed in the same way.

#### 2.4.1 Compared methods

We compare RPI-Zero with five state-of-the-art methods that (i) use an RNA/protein *sequence* as input, (ii) have public code to retrain the model with our benchmark data, and (iii) can input long transcripts. These models represent different strategies by using: (i) k-mer frequencies (IPMiner [27]); (ii) k-mer features and a graph of known interactions (NPI-GNN [31]); (iii) language-model embeddings and a graph neural network (ZHMolGraph [20]); (iv) fine-tuned language models joined by cross-attention (CORAL [5]); and (v) topological RNA features with an ESM-2 [19] protein embedding (ZHMolTopoRPI [7]). The last three were tested by their authors on molecules absent from training. Under our clustering rules, however, some of these test molecules are similar to training molecules (Supplementary Section 4.3). We retrain each method from scratch with its own features, architecture, and training procedure. We could not compare our model with BioLLMNet [1], RPIPLM [21], and PaRPI [29], because their RNA length limits fit only 14.6%, 3.8%, and 1.5% of the training examples, respectively. We also were unable to compare with the model of Yu et al. [48], which has no publicly released training code (Supplementary Section 3).

## 3 Results

### 3.1 Comparison with state-of-the-art methods

Table 2 compares RPI-Zero with the five state-of-the-art methods using the three test slices in our benchmark (unseen RNA, unseen protein, and unseen pair). Note that all models are trained from scratch on the training part of our benchmark. RPI-Zero has the best result of all eight metrics on all three slices. Its AUPRC is 2.7%, 7.4%, and 4.6% higher than that of the second-best method on unseen RNAs, proteins, and pairs. The second-best method changes from slice to slice. With respect to AUROC, it is ZHMolTopoRPI on unseen RNAs, IPMiner on unseen proteins, and CORAL on unseen pairs, and each of them ranks only third or fourth on other slices (Supplementary Fig. S1).

**Table 2.** Results on the three test slices. For the protein-macro (PM) metrics, we compute the metric on the examples of each protein with at least 10 positives and 10 negatives, and take the mean over these proteins. The other metrics are computed over all examples. Higher is better, and bold marks the best value of each slice.

| Method | PM-AUROC | PM-AUPRC | AUROC | AUPRC | Accuracy | Balanced acc. | F1 | MCC |
|---|---|---|---|---|---|---|---|---|
| *Unseen RNA* (177,838 examples, 180 proteins) |  |  |  |  |  |  |  |  |
| RPI-Zero | **0.758** | **0.681** | **0.769** | **0.753** | **0.691** | **0.693** | **0.689** | **0.385** |
| IPMiner | 0.711 | 0.626 | 0.731 | 0.713 | 0.664 | 0.657 | 0.604 | 0.323 |
| NPI-GNN | 0.465 | 0.405 | 0.594 | 0.564 | 0.586 | 0.575 | 0.472 | 0.161 |
| ZHMolGraph | 0.589 | 0.513 | 0.593 | 0.562 | 0.587 | 0.578 | 0.498 | 0.163 |
| CORAL | 0.662 | 0.565 | 0.720 | 0.704 | 0.656 | 0.649 | 0.593 | 0.307 |
| ZHMolTopoRPI | 0.729 | 0.643 | 0.756 | 0.733 | 0.686 | 0.682 | 0.650 | 0.367 |
| *Unseen protein* (269,408 examples, 24 proteins) |  |  |  |  |  |  |  |  |
| RPI-Zero | **0.708** | **0.606** | **0.724** | **0.637** | **0.660** | **0.664** | **0.642** | **0.326** |
| IPMiner | 0.675 | 0.579 | 0.666 | 0.593 | 0.624 | 0.587 | 0.414 | 0.214 |
| NPI-GNN | 0.535 | 0.471 | 0.568 | 0.510 | 0.580 | 0.544 | 0.357 | 0.106 |
| ZHMolGraph | 0.583 | 0.497 | 0.599 | 0.533 | 0.591 | 0.580 | 0.510 | 0.161 |
| CORAL | 0.635 | 0.532 | 0.650 | 0.560 | 0.608 | 0.568 | 0.371 | 0.174 |
| ZHMolTopoRPI | 0.640 | 0.539 | 0.641 | 0.551 | 0.609 | 0.608 | 0.570 | 0.214 |
| *Unseen pair* (29,808 examples, 23 proteins) |  |  |  |  |  |  |  |  |
| RPI-Zero | **0.680** | **0.578** | **0.680** | **0.589** | **0.634** | **0.631** | **0.589** | **0.260** |
| IPMiner | 0.608 | 0.503 | 0.605 | 0.512 | 0.580 | 0.523 | 0.185 | 0.080 |
| NPI-GNN | 0.462 | 0.386 | 0.534 | 0.483 | 0.570 | 0.500 | 0.000 | 0.000 |
| ZHMolGraph | 0.581 | 0.492 | 0.589 | 0.520 | 0.592 | 0.573 | 0.481 | 0.151 |
| CORAL | 0.638 | 0.542 | 0.657 | 0.563 | 0.614 | 0.571 | 0.371 | 0.182 |
| ZHMolTopoRPI | 0.609 | 0.513 | 0.616 | 0.529 | 0.591 | 0.584 | 0.531 | 0.168 |

We also compare our model with the methods in a pairwise manner for each protein. For each compared method (5) and each slice (3), we compute the AUPRC of RPI-Zero and of the compared method on the examples of each protein, and record the performance difference. Then, we take the mean over all tested proteins. We get a 95% confidence interval for this mean, by generating 10,000 bootstraps, each of which contains as many proteins as the slice. We statistically test the per-protein differences with a two-sided Wilcoxon signed-rank test [42] and correct the 15 *p*-values with the Holm method [13]. In 13 out of the 15 comparisons, RPI-Zero is significantly better than the compared method. In the two other cases, RPI-Zero is better than the compared method but the results are not statistically significant (corrected *p*-values are 0.24). These are IPMiner on unseen proteins and CORAL on unseen pairs. We also divide the test examples into groups by RNA type, RNA length, protein class, and protein length. RPI-Zero has the highest protein-macro AUROC in all 15 groups, and in 14 of them it is significantly better than each compared method (Supplementary Section 4.4).

We further investigate which models underperform when tested with novel molecules. The two methods that use a graph of known interactions are the weakest, because a new molecule has no edge in the training graph. NPI-GNN predicts every example of the unseen pair slice as negative. Protein language models do not close the gap either. CORAL and ZHMolTopoRPI describe the protein with ESM-2 [19]. Yet on unseen proteins, the protein-macro AUROC of RPI-Zero is 11.5% and 10.6% higher than theirs, and both of them are also below IPMiner, which uses only k-mer frequencies. RPI-Zero instead reads the protein and the RNA in one input and uses an RNA interaction foundation model, so the embedding of each nucleotide already depends on the protein. It is also the only method that learns from the position of each peak in the transcript, and the following sections show where this helps.

### 3.2 Separation of binding and non-binding pairs

AUROC and AUPRC depend only on the order (ranking) of the predictions, but a user who screens novel RNAs or proteins focuses on the probability of the interaction. We show that RPI-Zero separates the predicted probabilities of the two classes most clearly on every slice (Fig. 2). The gap between the median probabilities of its positive and its negative examples is 0.349 on unseen RNAs, 0.207 on unseen proteins, and 0.156 on unseen pairs, against at most 0.198, 0.110, and 0.091 for other compared methods. RPI-Zero has a large variance; 80%, 72%, and 71% of its predictions are not within the 0.4–0.6 range, for each respective slice. We observe that while ZHMolTopoRPI is the second-best method by its mean rank in Table 2 (also see Supplementary Fig. S1), its variance is very low; on unseen proteins and unseen pairs 90% of its predictions lie between 0.45 and 0.53. The medians of its positive and its negative examples differ by only 0.012 and 0.009. On unseen pairs, NPI-GNN gives every example a probability between 0.359 and 0.360, which appears as a line in Fig. 2. Neither molecule of these pairs has an edge in the training graph. Both get a zero node2vec vector and no neighbors, so only the k-mer frequencies differ between pairs (Supplementary Section 3.2).

**Fig. 2.**
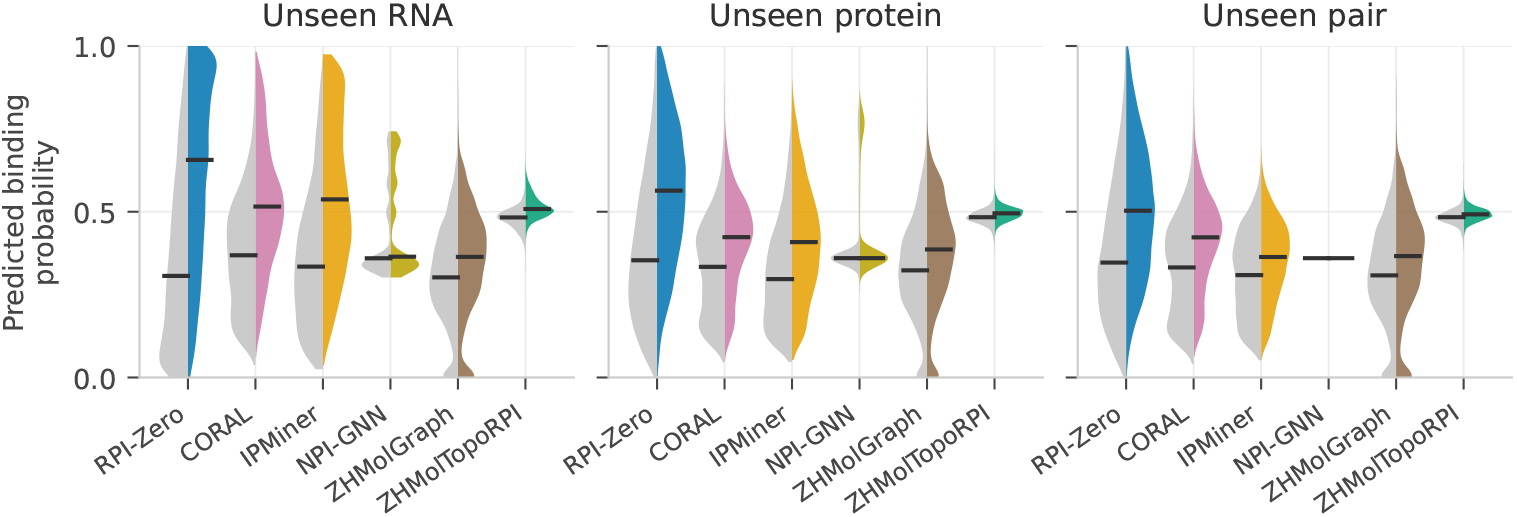
Predicted binding probabilities on the three test slices. The left half of each violin shows the negatives and the right half the positives, each from up to 20,000 random examples. Black lines mark the medians.

The difference is largest for new proteins. On unseen proteins, RPI-Zero gives a probability of at least 0.6 to 44% of the binding pairs, against 15%, 7%, and 0% for IPMiner, CORAL, and ZHMolTopoRPI, respectively, which are the next best methods with respect to AUROC. On unseen pairs, this share is 33% for RPI-Zero and at most 7% for the compared methods. Of the pairs with a probability of at least 0.6, 67% bind on unseen proteins and 63% on unseen pairs, while only 44% and 43% of all pairs bind. At a precision of 60%, RPI-Zero detects 68% of the binding pairs on unseen proteins and 46% on unseen pairs, against at most 47% and 37% for other compared methods.

### 3.3 Detection of sparse binding signals

A protein often binds only a short stretch of a long RNA. Between 43% and 49% of the positive examples of the test slices hold a single whole peak bound by the protein. This peak covers a median of 25 to 32 nt, which is 1.2% to 1.7% of the RNA sequence in that pair. A method that summarizes the whole RNA in one vector or in k-mer counts mixes this short site with thousands of unbound nucleotides.

Fig. 3a groups the positive examples by the number of whole peaks on their RNA that are bound by their protein. For each protein, we compute the AUROC of a method on the positive examples of one group and all negative examples of that protein, so only the positive examples change from group to group.

**Fig. 3.**
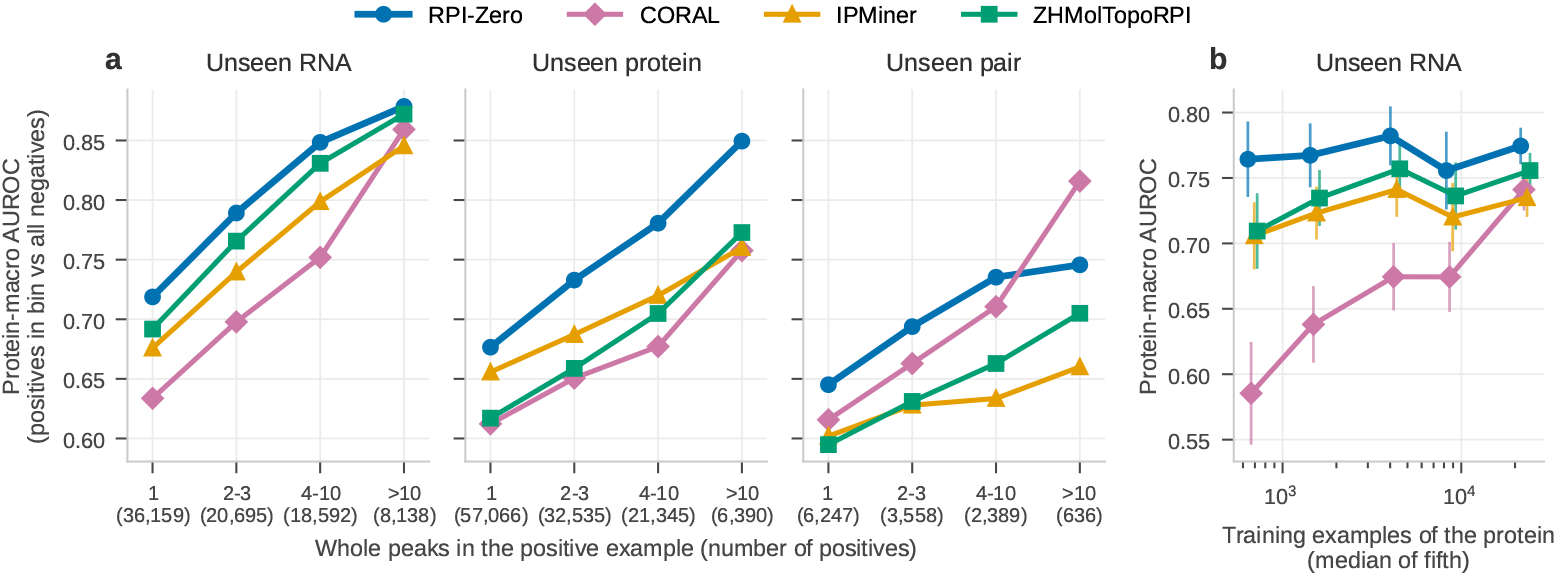
(a) Protein-macro AUROC for positive examples grouped by the number of whole peaks on their RNA that are bound by their protein. The mean is over the proteins with at least 10 positive examples in the group and 10 negative examples, and the number of positives of each group is given below the axis. (b) Protein-macro AUROC on the unseen RNA slice against the number of training examples of the protein. The 157 proteins of the slice that are in the training data are divided into five groups of equal size by their number of training examples. Each point is the protein-macro AUROC over the proteins of a group, placed at the median number of training examples of the group and shifted slightly sideways for readability. Error bars are 95% confidence intervals of the mean. NPI-GNN and ZHMolGraph are not shown. They stay below 0.62 in every group of (a), and below 0.50 and 0.63, respectively, in every group of (b).

The single-peak group is the hardest group, and it is also the largest one. RPI-Zero is the best method on this group on all three slices, with a protein-macro AUROC of 0.719, 0.677, and 0.645. The second-best method is different on each slice, with 0.692 for ZHMolTopoRPI, 0.656 for IPMiner, and 0.616 for CORAL. On unseen RNAs and unseen proteins, RPI-Zero is the best method in all four groups. On unseen proteins, its lead grows with the number of peaks.

On unseen RNAs, the AUROC of CORAL falls substantially (™0.226) in detecting interactions with a single peak compared to the examples with more than 10 peaks. CORAL is ahead of RPI-Zero only in the group with more than 10 peaks on unseen pairs, which holds 5% of the positive examples of the slice.

### 3.4 Proteins with scarce training data

Most human RNA-binding proteins have little or no CLIP data. Fig. 3b shows that the accuracy of RPI-Zero on unseen RNAs does not depend on how much training data a protein has.

The Spearman correlation between the AUROC of a method on a protein and the number of training examples of that protein is 0.036 for RPI-Zero, the lowest of the six methods, against 0.148 for IPMiner, 0.208 for ZHMolTopoRPI, and 0.546 for CORAL. RPI-Zero reaches 0.761 on the third of the proteins with the fewest training examples (218 to 1,702) and 0.764 on the third with the most (7,191 to 32,238), while CORAL falls from 0.720 on the proteins with the most data to 0.604 on those with the fewest. The lead of RPI-Zero is therefore largest where data is scarce. On the proteins with the fewest and with the most examples, its lead is 0.047 and 0.018 over ZHMolTopoRPI, 0.052 and 0.037 over IPMiner, and 0.157 and 0.044 over CORAL. Two parts of the design may help here. The sampler gives a protein with few examples more draws than its share of the data (Section 2.3). All proteins also share the same 16 motif prototypes (Section 2.2), so a protein with few examples is described by the same vectors as the proteins with many examples.

### 3.5 Localization of binding sites

The five compared methods give one score for an RNA-protein pair. RPI-Zero produces a logit for every nucleotide (Section 2.2), so each prediction comes with a profile across the RNA sequence that marks the likely binding sites. Fig. 4a shows two examples with proteins absent from training: GEMIN5 interacts with a chunk of the UBR5 mRNA (unseen pair), and GTF2F1 interacts with the MEX3D mRNA (unseen protein, seen RNA). Each RNA molecule holds a single CLIP peak of about 100 nt, and the highest logit the model produces lies in this peak. The AUPRC of the logits produced is 0.84 and 0.81, respectively. A random ranking would achieve an AUPRC of 0.04 and 0.03, respectively.

**Fig. 4.**
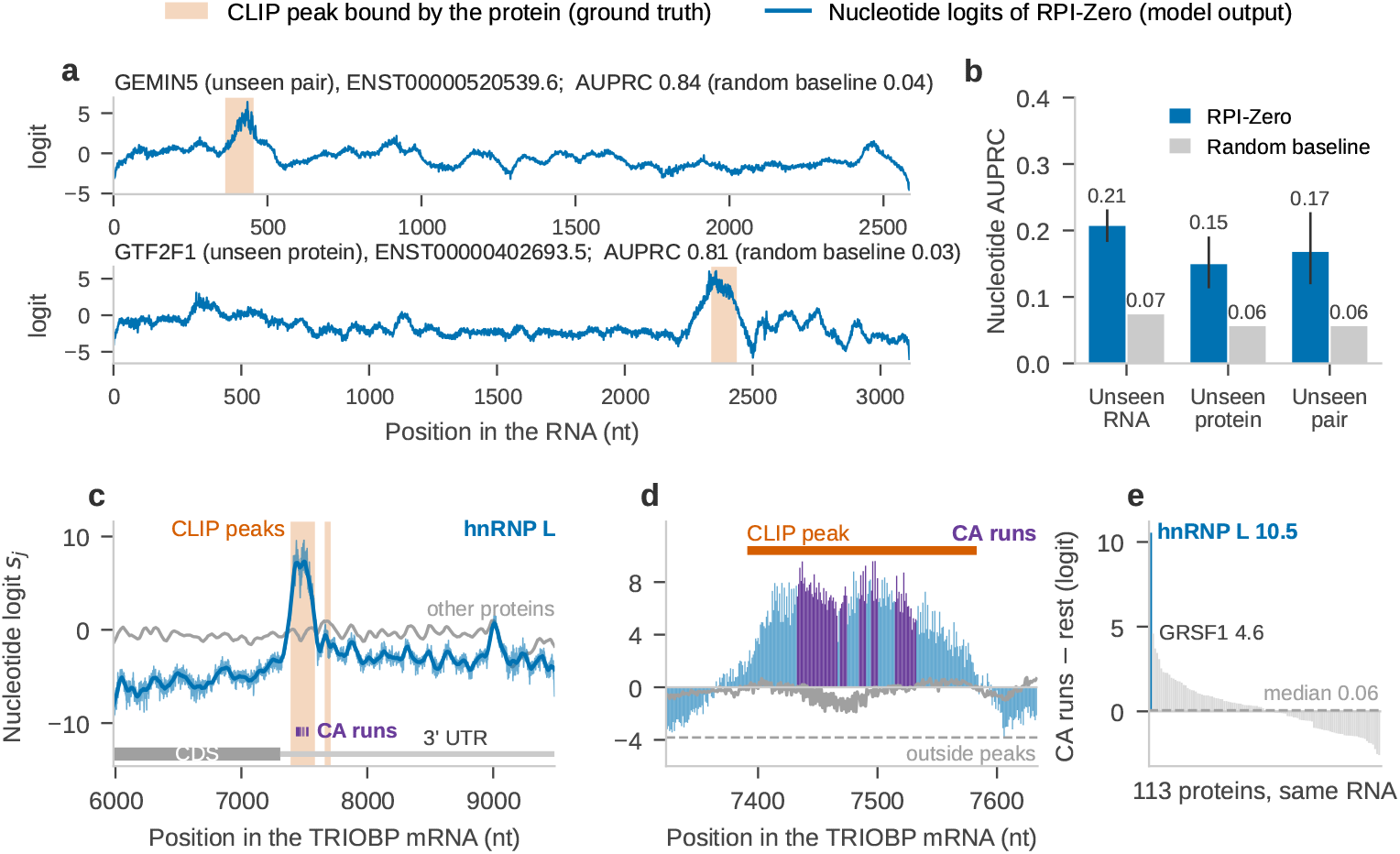
Nucleotide logits of RPI-Zero. (a) Nucleotide logits *s*_*j*_ of RPI-Zero (blue) along the RNA of two positive examples with proteins absent from training, GEMIN5 (unseen pair) and GTF2F1 (unseen protein). The ground-truth CLIP peak bound by the protein is orange, and titles give the AUPRC of the logits with its random baseline. (b) AUPRC of the nucleotide logits within each positive example and its random baseline, the share of the nucleotides in a peak, averaged over the examples of each protein and then over the proteins. We use 4,000 random positive examples per test slice, and error bars are 95% bootstrap confidence intervals over proteins. (c–e) hnRNP L on a chunk of the TRIOBP mRNA from the unseen RNA slice. (c) Logits (thin blue line), their 41-nt window mean (thick blue line), eCLIP peaks (orange), and runs of three or more CA dinucleotides (purple). (d) Logits around the larger peak, with CA-run nucleotides in purple. The dashed line is the mean logit outside the peaks, and grey lines in (c) and (d) are the median logits of the 112 other proteins of (e). (e) Mean logit over the CA runs minus that over the rest of the chunk, for 113 proteins.

Inspired by these examples, we further test whether the high logits fall into the CLIP peaks across our test data. We compare logits only within each positive example, so the result does not depend on how the example ranks against other examples. CLIP peaks cover only 5.6% to 7.4% of the RNA sequence in a positive example, and a random ranking achieves an AUPRC equal to this percentage (Fig. 4b). The AUPRC of the nucleotide logits produced by our model is 3.2 times higher than this random baseline in unseen RNAs, 2.7 times higher in unseen proteins, and 3.2 times higher in unseen pairs. It is higher than the random baseline for all proteins in all three slices. We also take the 41-nt window with the highest mean logit in each example. This is the window of the pooling, and its width is close to the length of a CLIP peak (Section 2.2). This window overlaps a CLIP peak 2.6, 2.2, and 2.5 times more than a window placed at random in the same RNA sequence. The ratio ranges from 2.6 to 2.8 times when all peaks lie at least 300 nt from both ends, so the logits do not fix on the ends of the RNA sequence.

The gain over the random baseline is even larger when the binding is sparse. In 30% to 43% of the positive examples, the peaks cover less than 2% of the RNA. On these examples, the AUPRC of the nucleotide logits is 7.2, 4.9, and 5.8 times the random baseline, and the highest window overlaps a peak 4.5, 2.8, and 3.2 times more than a random window. The pair logit of RPI-Zero is pooled from the nucleotide logits, so such an RNA sequence gets a high score when the logits find its short binding site (Section 3.3). RPI-Zero therefore localizes binding for proteins and RNAs that it has never seen.

### 3.6 Recognition of a known binding motif

Fig. 4c–d shows a positive example that only RPI-Zero predicts as binding. The hnRNP L protein interacts with a 3,504-nt chunk of the TRIOBP mRNA from the unseen RNA slice. A chunk is a part of the transcript, because the entire 10,085-nt mRNA does not fit in the model together with hnRNP L (Section 2.1). The probability assigned by RPI-Zero is 0.996, while the five compared methods yield 0.18 to 0.45, below their decision thresholds. RPI-Zero also places this pair in the top 5% of the examples of hnRNP L in this test slice, while every compared method places it below the median. Of the six methods, only RPI-Zero gives a logit for every nucleotide (Section 3.5), so only its prediction shows which part of the chunk is bound. The larger eCLIP peak of the chunk [38] lies in the 3^′^ UTR and holds all runs of three or more CA dinucleotides in the chunk. In vitro selection (SELEX) showed that hnRNP L binds CA repeats and CA-rich elements [14], and its binding sites in cells are rich in C and A [30]. The logits of RPI-Zero rise over the CA runs (Fig. 4c,d). The nucleotides of these runs have a mean logit of 7.1, against 3.3 for the rest of the peak and ™3.8 outside the peaks, and the highest logit of the chunk lies in a CA run.

This rise is specific to the protein. We scored this chunk with the 113 proteins that fit into one input with it, hnRNP L included. The mean logit over the CA runs minus that over the rest of the chunk is 10.5 for hnRNP L, 4.6 for the next protein, and 0.06 for the median protein (Fig. 4e). In the 46 positive examples of hnRNP L in this slice that contain a CA run, the logits rank CA-run nucleotides above the others with a median AUROC of 0.91.

## 4 Discussion

We presented a benchmark and a model for RNA–protein interaction prediction on molecules absent from training. On this benchmark, the second-best method is different on each test slice. The two graph methods are the weakest on new molecules, and the ESM-2 protein embedding does not bring CORAL or ZHMolTopoRPI above IPMiner on unseen proteins. Results on one kind of held-out molecule therefore do not show how a method works on another kind. RPI-Zero has the best results of all eight metrics on all three slices.

We see three reasons for these results. First, RNA-X reads the protein and the RNA in one input, so the logit of each nucleotide depends on the protein (Section 3.6). Second, the model is trained on full transcripts with a label at each nucleotide, and the pair score is pooled from the nucleotide logits. It is the best method on positives with a single peak (Section 3.3), and it localizes the peaks for proteins and RNAs absent from training (Section 3.5). Removing the nucleotide loss, the pair loss, or the convolution blocks lowers the accuracy on all three validation slices (Supplementary Section 4.2). Third, one model is trained on all proteins with a balanced sampler, and its accuracy on unseen RNAs does not depend on how much CLIP data a protein has (Section 3.4). Its probabilities also separate the two classes more clearly than those of the compared methods (Section 3.2), so the probability itself can be used to screen new molecules. A new RNA or protein needs only its sequence, with no structure and no known interactions. Training updates only the head of 2.0 million parameters and takes 12 hours on one GPU.

The benchmark and the model have limits that follow from the available data. The main one is the number of proteins. CLIP data exist for about 10% of human RNA-binding proteins [48], and the 209 proteins of the benchmark form 181 clusters at 30% identity. Like the stronger compared methods, RPI-Zero is less accurate on unseen proteins and unseen pairs than on unseen RNAs. Because these slices hold 24 proteins each, we compare the methods protein by protein with bootstrap intervals and corrected tests (Section 3.1). Proteins from new CLIP experiments can be added by retraining only the head. The labels follow the limits of CLIP. CLIP finds binding sites only on RNAs expressed in the profiled cells, so some negatives can be bound in other cells. We reduce such cases by calling a transcript unbound only if no peak bound by the protein lies on its genomic span in the four databases. All methods are tested on the same labels. The model has no input for the cell type, so it predicts whether a protein can bind an RNA and not whether it binds it in a given cell.

Two limits come from the input. The benchmark labels binding in the exons of one transcript per gene, because a peak is kept only if it lies fully inside one exon. Every label is then exact on the spliced transcript, but binding in introns, across exon junctions, and on other isoforms is not labeled. The same chunking applies to pre-mRNA sequences, so the benchmark can be extended to intronic binding. The input of RNA-X has 4,096 tokens, so a long transcript is scored chunk by chunk, and combining the chunk scores into one score per transcript is left for future work.

