## Supplementary Material for "RPI-Zero: Zero-shot RNA-Protein Interaction Prediction"

### 1 Dataset

This section gives the details of the benchmark in Section 2.1 of the main text, in the same order. Every random choice in the construction uses a fixed seed or a fixed hash, so the benchmark can be rebuilt exactly. The benchmark is available at https://zenodo.org/records/23748141. The code that builds the benchmark, trains RPI-Zero, and evaluates it is available at https://github.com/ciceklab/RPI-Zero, together with the trained model. Table S1 gives the counts from the peaks to the examples, and Table S2 gives the size of each slice.

#### 1.1 CLIP peaks and proteins

We take the CLIP peaks of CLIPdb [30], ENCODE eCLIP [27], POSTAR3 [33], and POSTAR2 [36]. Each CLIP experiment targets one protein. A peak is an RNA region that this protein binds, given as an interval on the hg38 genome. The benchmark has 11,607,227 peaks bound by 209 proteins. The number of peaks bound by one protein ranges from 85 to 1,079,118, with a median of 11,628. We describe each protein by its amino acid sequence, which has 127 to 2,804 amino acids, with a median of 577. The databases report only binding, so we build the negative examples ourselves (Section 1.5).

#### 1.2 Transcripts

We use the GENCODE release 50 annotation [19] and the hg38 genome. There are 29,989 genes with at least one peak that lies fully inside an exon of one of their transcripts. For each of these genes, we choose one transcript among the transcripts that hold such a peak. We prefer the MANE Select transcript [18], then the Ensembl canonical transcript [31], and then the longest transcript. This gives 16,801 MANE Select, 11,751 Ensembl canonical, and 1,437 longest transcripts. The choice depends only on the gene, so all proteins see the same transcript of a gene. We join the exons in transcript order on the strand of the gene. The median transcript has 2,023 nt.

We map a peak to a chosen transcript if the peak overlaps the genomic span of the transcript on the same strand. We keep the peak only if it lies fully inside one exon. We do not clip a peak that crosses an exon border, because the clipped peak would end at the exon border, which is not where the binding site ends.

#### 1.3 Examples and labels

The input of RNA-X [25] has at most 4,096 tokens, and the protein, the RNA, and three special tokens share it. For a protein of $L_p$ amino acids, the RNA can therefore have at most $4{,}093 - L_p$ nt. We call this length the budget of the protein. The budget is 3,966 nt for the shortest protein and 1,289 nt for the longest. Of the transcripts, 23% are longer than the largest budget and 64% are longer than the smallest budget.

We cut a transcript that is longer than the budget into parts on a fixed grid. The first part starts at the $5'$ end, and each next part starts one stride later. The stride is the budget minus 512 nt, so neighboring parts overlap by 512 nt. Every peak shorter than 512 nt therefore lies whole in at least one part. The grid does not depend on the peaks, so the position of a peak inside a part does not reveal the label.

Before we label, we merge the overlapping peaks that one protein binds on a transcript. An example is a protein paired with a transcript or with a part of one. We make one example for each pair of a part and a protein, and not one for each peak. With one example for each peak, the same pair would appear several times with different partial labels. The nucleotides inside a whole peak have the label 1, and all other nucleotides have the label 0. The only exception is the piece of a peak that a part border cuts. We mask this piece, because the part does not show the whole site. An example is positive if its RNA holds at least one whole peak bound by the protein. In the training slice, whole peaks cover a median of 4.3% of the RNA of a positive example, so the binding signal is sparse.

#### 1.4 Clustering and data splits

We split the RNAs and the proteins by clusters of sequence similarity, and we never divide a cluster between buckets.

**1.4.1 RNA axis:** We cluster whole transcripts and not peak sequences, because the model reads whole transcripts. Two peaks on the same transcript can fall into different clusters of peak sequences, so clusters of peak sequences do not control the similarity between transcripts. We cluster the 29,989 transcripts with MMseqs2 [26] at a minimum sequence identity of 0.8 and a coverage of 0.8 of the target sequence. With this coverage, a short transcript that lies inside a long paralog joins the cluster of the paralog. This gives 26,684 clusters. We shuffle the clusters and fill four groups with whole clusters up to the shares 0.78, 0.02, 0.10, and 0.10. The first two groups form the training bucket, and the other two form the validation and the test bucket. We do not train on the transcripts of the second group, which we call the monitor transcripts (Section 1.4.4). We count these shares in examples and not in transcripts, because the number of examples per transcript varies. The realized shares are 0.7802, 0.0199, 0.0999, and 0.1000, so the training bucket holds 80% of the examples.

**1.4.2 Protein axis:** We cluster the 209 proteins with MMseqs2 [26] at a minimum sequence identity of 0.3 and a sensitivity of 7.5, which gives 181 clusters. A deterministic mixed-integer program assigns whole clusters to the training, validation, and test buckets. Its target is that the proteins of the three buckets bind 80%, 10%, and 10% of the peaks. The result is 80.4%, 9.8%, and 9.8% of the peaks, with 161, 24, and 24 proteins.

**1.4.3 Slices:** The RNA bucket and the protein bucket of an example give its slice (Table 1 of the main text). We select models on the validation slices, so no test molecule may appear in them. A pair of a test RNA and a validation protein therefore goes to the test unseen RNA slice, and a pair of a validation RNA and a test protein goes to the test unseen protein slice. Both molecules of these pairs are absent from the training data.

**1.4.4 Monitor slice:** The 637 monitor transcripts and the training proteins give the monitor slice, with 30,295 examples (Table S2). We score this slice during training to follow the progress of training. We do not train on it, and we do not use it for model selection or testing. Table 1 of the main text therefore leaves it out. With a validation or a test protein, a monitor transcript goes to an unseen protein slice, like every other transcript of the training bucket. This gives 5,535 examples of the validation unseen protein slice and 5,925 examples of the test unseen protein slice.

#### 1.5 Negative examples

A negative example is an example whose RNA holds no whole peak bound by the protein. We build two kinds of negatives.

**1.5.1 Same-transcript negatives:** These negatives come from transcripts that the protein binds. A long transcript has many parts that touch no peak bound by the protein. For each protein and each transcript that holds a peak bound by it, we rank these parts by a hash of the transcript, the protein, and the index of the part, and we keep the first two.

**1.5.2 Cross-transcript negatives:** These negatives come from transcripts that the protein does not bind. A transcript is unbound for a protein if no peak bound by the protein lies on the genomic span of the transcript on the same strand, introns included. This check uses all peaks bound by the protein in the four databases, including the peaks that do not lie fully inside an exon. No known binding site of the protein therefore lies on the genomic span of a cross-transcript negative. We draw the negatives for the positives of a protein in one RNA bucket from the unbound transcripts of the same bucket. A negative is therefore in the same slice as its positives, and no negative crosses a split. We take at most one negative for each positive and at most two parts from each unbound transcript.

**1.5.3 Matching:** We match two properties of the cross-transcript negatives to those of the positives, because a model could read either property without looking at the protein. The first property is the length of the RNA. A part that fills the budget has the length $4{,}093 - L_p$, so its length reveals the protein. The second property is the partner count of the transcript, which is the number of other proteins that bind at least one peak on its genomic span. A transcript that many proteins bind is a positive for many proteins and is rarely unbound. Without matching, the negatives would therefore have fewer partners than the positives.

We match the two properties jointly, because long transcripts collect more peaks and therefore more partners. For each protein and RNA bucket, we divide the positive examples into at most 32 bins by the length of the RNA and at most 8 bins by the partner count, with equal numbers of positives in each bin. Each combination of a length bin and a partner bin asks for as many negatives as it holds positives. We go through the unbound transcripts in a fixed pseudo-random order and fill each combination with parts from the same length bin and the same partner bin. A combination that cannot be filled in this way gets parts from the same length bin and the nearest partner bin, and we take these parts from the transcripts with the most partners first. We use this second pass instead of leaving the combination empty, because an empty combination would lower the share of negatives of the protein. The benchmark has 797,774 cross-transcript negatives.

#### 1.6 Checks on the final data

We run four checks on all 2,074,753 examples, which are the checks of Fig. 1f of the main text. First, no data leaks between buckets. Each transcript lies in only one RNA bucket and each protein in only one protein bucket, and no transcript is both a positive and a cross-transcript negative of the same protein. Second, the labels agree with the peaks. An example is positive exactly when its RNA holds a whole peak bound by the protein, and every labeled peak and every masked piece lies inside the RNA of the example without overlap. Third, every example fits into the input of RNA-X. No input is longer than 4,096 tokens, and no part is longer than the budget of its protein. Fourth, the centers of the whole peaks are spread over the whole length of the parts. They do not gather at the center of a part, as they would if the parts were placed around the peaks.

#### 1.7 Statistics of the benchmark

In the training slice, 48.1% of the examples are positive, 14.2% are same-transcript negatives, and 37.7% are cross-transcript negatives. Over all slices, these shares are 46.8%, 14.7%, and 38.5%.

**Table S1.** From peaks to examples.

| Step | Count |
|---|---|
| CLIP peaks | 11,607,227 |
| Genes with a peak fully inside an exon (one transcript each) | 29,989 |
| Pairs of a transcript and a protein with at least one peak inside an exon | 732,161 |
| Positive examples | 971,450 |
| Same-transcript negatives | 305,529 |
| Cross-transcript negatives | 797,774 |
| All examples | 2,074,753 |

**Table S2.** Size of each slice. A positive example holds at least one whole peak bound by the protein. Peak nt is the share of the labeled nucleotides of a slice that lie inside a peak. The monitor slice is used only to follow training (Section 1.4.4).

| Slice | Examples | Positives | Transcripts | Proteins | Peak nt (%) |
|---|---|---|---|---|---|
| Training | 1,169,793 | 562,637 | 23,196 | 161 | 3.93 |
| Monitor | 30,295 | 14,586 | 637 | 160 | 3.74 |
| Validation, unseen RNA | 149,016 | 71,906 | 3,004 | 161 | 3.85 |
| Validation, unseen protein | 221,288 | 96,668 | 20,572 | 24 | 2.16 |
| Validation, unseen pair | 27,307 | 11,903 | 2,567 | 23 | 2.06 |
| Test, unseen RNA | 177,838 | 83,584 | 3,149 | 184 | 3.52 |
| Test, unseen protein | 269,408 | 117,336 | 24,099 | 24 | 1.90 |
| Test, unseen pair | 29,808 | 12,830 | 2,755 | 24 | 1.86 |

### 2 Architecture and training of RPI-Zero

This section gives the details of Sections 2.2 and 2.3 of the main text. We describe the model for a single example, and we omit the bias terms of the linear layers from the equations. Table S3 lists the hyperparameters.

#### 2.1 Input and encoder

The input is one sequence of tokens. It holds a `[CLS]` token, the protein, the separator `$`, the RNA, and an `[EOS]` token. RNA-X [25] has separate tokens for amino acids and for nucleotides, and its vocabulary has 36 tokens. The embedding of a token is the sum of a token embedding, a learned position embedding, and a residue type embedding. The residue type marks whether the token is an amino acid, a nucleotide, or a special token. The protein is always given in full, and the RNA is a transcript or a part of one (Section 1.3).

RNA-X is a transformer [28] with 14 layers, a width of 768, 12 attention heads, and 136 million parameters. It was pretrained on more than 100 million interactions of RNA with proteins, RNA, and DNA [25]. Self-attention runs over the whole input. The embedding of each nucleotide therefore depends on the protein, and the embedding of each amino acid depends on the RNA. For this reason, the head needs no separate protein encoder and no cross-attention between the two molecules. We keep all weights of RNA-X frozen. We take the output of its last layer before the final layer normalization and split it at the separator. This gives the protein embeddings $E^p \in \mathbb{R}^{L_p \times 768}$ and the RNA embeddings $E^r \in \mathbb{R}^{L_r \times 768}$, where $L_r$ is the length of the RNA. The special tokens are dropped.

#### 2.2 Motif prototypes

The head works at a width of $d = 256$ and uses a dropout rate of 0.1. It averages the rows of $E^p$ into a vector $\bar{\mathbf{e}}^p \in \mathbb{R}^{768}$ and computes the protein vector $\mathbf{p} = \mathrm{GELU}(W_{\mathrm{prot}}\, \bar{\mathbf{e}}^p) \in \mathbb{R}^d$ [10]. The head holds $K = 16$ motif

**Table S3.** Hyperparameters of RPI-Zero.

| Hyperparameter | Value | Hyperparameter | Value |
|---|---|---|---|
| *Head* | | *Optimization* | |
| Width of the head, $d$ | 256 | Optimizer | AdamW |
| Motif prototypes, $K$ | 16 | Learning rate | $5 \times 10^{-4}$ |
| Convolution blocks | 5 | Weight decay | 0.01 |
| Kernel width of the convolutions | 129 | Warmup | 5% of the steps |
| Receptive field of $\mathbf{h}_j$ | 641 nt | Schedule after warmup | cosine decay to 0 |
| Window of the mean before pooling | 41 nt | Gradient norm clipping | 1.0 |
| Initial pooling temperature $\tau$ | 1.0 | Training steps | 154,000 |
| Dropout | 0.1 | *Batch sampling* | |
| Trainable parameters | 2,019,862 | Maximum tokens per batch | 65,536 |
| *Loss* | | Maximum examples per batch | 32 |
| Weight of positive nucleotides, $w_+$ | 49.06 | Exponent of the protein draws, $n_p^e$ | $e = 1/2$ |
| Steps with $\mathcal{L}_{\text{nt}}$ only, $T_0$ | 15,400 | Share of positives in the draws of a protein | 0.5 |
| Steps of the $\lambda$ ramp, $T_1$ | 15,400 | *Model selection* | |
| Final weight of $\mathcal{L}_{\text{pair}}$, $\lambda_{\max}$ | 10 | Interval between validation checks | 10,000 steps |
| Examples used to set $w$ and $b$ | 2,048 | Decision threshold | 0.439 |

prototypes $\mu_1, \ldots, \mu_K \in \mathbb{R}^d$, which are learned vectors that all proteins share. A router gives the weight of each prototype for the protein, and the weighted sum of the prototypes is the protein context $\mathbf{q}$,

$$\mathbf{a} = \operatorname{softmax}(W_{\text{route}}\, \mathbf{p}), \qquad \mathbf{q} = \sum_{k=1}^{K} a_k\, \mu_k. \tag{S1}$$

The prototypes are trained with the rest of the head and are not fixed to known motifs. Because $\mathbf{q}$ is a weighted mean of the shared prototypes, a protein that is absent from training is described by the same 16 vectors as the training proteins. Only its weights over these vectors are new. The weights $a_k$ are used twice. They form $\mathbf{q}$, which gates the convolutions over the RNA (Eq. (S2)), and they weight the motif score of each nucleotide (Eq. (S3)). The router starts with zero weights, so at the start of training every protein gives the same weight to all prototypes.

#### 2.3 Protein-gated convolutions

The head maps each row $\mathbf{e}_j^r$ of $E^r$ to width $d$ with a linear layer, a layer normalization, and a GELU. A position-wise layer follows, with a linear map, a GELU, dropout, and a layer normalization. It gives the representation $\mathbf{h}_j^{(0)}$ of nucleotide $j$. Five residual blocks then mix the representations along the RNA. Block $\ell$ computes

$$\begin{aligned} [\mathbf{v}_j;\, \mathbf{u}_j] &= W_{\text{in}}^{(\ell)}\, \mathrm{LN}\big(\mathbf{h}_j^{(\ell-1)}\big), \\ \tilde{\mathbf{v}}_j &= \sum_{o=-64}^{64} \boldsymbol{\kappa}_o^{(\ell)} \odot \mathbf{v}_{j+o}, \\ \mathbf{h}_j^{(\ell)} &= \mathbf{h}_j^{(\ell-1)} + W_{\text{out}}^{(\ell)} \Big[\mathrm{GELU}(\tilde{\mathbf{v}}_j) \odot \sigma\big(\mathbf{u}_j + W_{\text{gate}}^{(\ell)}\, \mathbf{q}\big)\Big], \end{aligned} \tag{S2}$$

where $\mathbf{v}_j, \mathbf{u}_j \in \mathbb{R}^d$ are the two halves of a linear map of width $2d$, LN is layer normalization, $\odot$ is the elementwise product, $\sigma$ is the sigmoid function, and $\mathbf{v}_{j+o} = \mathbf{0}$ when $j + o$ lies outside the RNA.

The second line of Eq. (S2) is a depthwise convolution. Each of the $d$ channels has its own kernel of width 129, and $\boldsymbol{\kappa}_o^{(\ell)} \in \mathbb{R}^d$ holds the weights of all channels at offset $o$. The convolution mixes positions but not channels, and $W_{\text{in}}^{(\ell)}$ and $W_{\text{out}}^{(\ell)}$ mix channels at each position. As in gated convolutional networks [6], a sigmoid gate sets how much of the convolution output enters the residual sum, for each channel and each nucleotide.

The gate depends on the nucleotide through $\mathbf{u}_j$ and on the protein through $\mathbf{q}$, so the same RNA can give different features for different proteins. Dropout is applied before $W_{\text{out}}^{(\ell)}$. The matrix $W_{\text{out}}^{(\ell)}$ starts at zero, so each block passes its input unchanged at the start of training. Each block widens the receptive field by 128 nt. After five blocks, the representation $\mathbf{h}_j = \mathbf{h}_j^{(5)}$ depends on the RNA embeddings of the 320 nucleotides on each side of $j$, which is a window of 641 nt.

#### 2.4 Nucleotide logits

The logit of nucleotide $j$ is the sum of four terms,

$$s_j = \mathbf{w}_s^\top \mathbf{h}_j + \sum_{k=1}^{K} a_k \frac{\langle W\mathbf{h}_j,\, \mu_k \rangle}{\sqrt{d}} + \mathbf{w}_p^\top \mathbf{p} + \mathbf{w}_r^\top \bar{\mathbf{h}}, \tag{S3}$$

where $W$ is a learned $d \times d$ matrix and $\bar{\mathbf{h}} = \frac{1}{L_r} \sum_{j=1}^{L_r} \mathbf{h}_j$ is the mean representation of the RNA. The position term $\mathbf{w}_s^\top \mathbf{h}_j$ reads the representation of the nucleotide, which depends on the protein through RNA-X and through the gates. The motif score maps the representation with $W$, compares it with each prototype by a scaled dot product, and weights the comparisons with the weights $a_k$ of the protein. A nucleotide therefore gets a high motif score when its representation matches the prototypes that the protein selects. The protein term depends only on the protein vector $\mathbf{p}$, and the RNA term depends only on the mean representation $\bar{\mathbf{h}}$. Each of these two terms adds the same value to all nucleotide logits of an example, and both terms start at zero. We train the nucleotide logits against the peak labels with $\mathcal{L}_{\text{nt}}$ (Eq. (S7)).

#### 2.5 Pooling

The pooling turns the nucleotide logits into one score $z$ for the pair. It first averages the logits over a window of 41 nt centered on each nucleotide,

$$\bar{s}_j = \frac{1}{|\Omega_j|} \sum_{j' \in \Omega_j} s_{j'}, \qquad \Omega_j = \{\, j' : |j' - j| \le 20,\ 1 \le j' \le L_r \,\}, \tag{S4}$$

and then applies a log-sum-exp with a learned temperature $\tau$ and a correction for the length of the RNA,

$$z = \frac{1}{\tau} \Big( \log \sum_{j=1}^{L_r} e^{\tau \bar{s}_j} - \log L_r \Big). \tag{S5}$$

The pair logit is $g = w\,z + b$, where $w$ and $b$ are two learned scalars that we set once at the end of the first training phase (Section 2.6). The binding probability of the pair is $\sigma(g)$.

Equation (S5) is $1/\tau$ times the logarithm of the mean of $e^{\tau \bar{s}_j}$. The score $z$ therefore lies between the mean and the maximum of the windowed logits. It approaches the mean as $\tau \to 0$ and the maximum as $\tau \to \infty$. Every nucleotide contributes to $z$, and the highest windowed logits contribute the most. The temperature starts at 1 and is 1.64 at the selected checkpoint.

The term $-\log L_r$ removes the effect of the length of the RNA. If every $\bar{s}_j$ equals the same value $c$, then $z = c$ for any $L_r$. Without this term, a longer RNA would get a higher score only because the sum has more terms. The length of a part is $4{,}093 - L_p$ and depends on the protein, so without this term the pair logit could partly encode the length of the protein instead of binding.

The log-sum-exp does not depend on the order of the nucleotides. Without the window mean, 41 high logits scattered along the RNA would give the same score as 41 high logits in a row. A CLIP peak is a run of adjacent nucleotides, with a median length of 35 nt in our data. The window mean divides the excess of a single high logit over its neighbors by 41, and it keeps the value of a run that fills the window. The pair logit therefore responds to stretches of high logits more than to single nucleotides. The window mean is used only in the pooling, so $\mathcal{L}_{\text{nt}}$ is computed on the logits $s_j$ themselves.

The pooling uses all nucleotides of the RNA, including the masked pieces of cut peaks, which are left out of $\mathcal{L}_{\text{nt}}$ only. Padding is left out of every step that mixes positions. These steps are the convolutions, the mean $\bar{\mathbf{h}}$, the window mean, and the log-sum-exp.

#### 2.6 Training objective

We train only the head, which has 2.0 million parameters, and keep all weights of RNA-X frozen. For example $i$, the head gives a logit $s_{ij}$ for each nucleotide $j$ of the RNA and a pair logit $g_i$. The loss has a nucleotide term and a pair term,

$$\mathcal{L} = \mathcal{L}_{\text{nt}} + \lambda(t)\,\mathcal{L}_{\text{pair}}, \tag{S6}$$

where $t$ is the optimizer step. The nucleotide term is a weighted binary cross-entropy over the nucleotides that carry a label,

$$\mathcal{L}_{\text{nt}} = -\frac{1}{\sum_{i,j} m_{ij}} \sum_{i,j} m_{ij} \Big[ w_+ \, y_{ij} \log \sigma(s_{ij}) + (1 - y_{ij}) \log\big(1 - \sigma(s_{ij})\big) \Big], \tag{S7}$$

where $y_{ij}$ is the label of the nucleotide, and $m_{ij}$ is 0 for padding and for the masked piece of a cut peak and 1 otherwise. Only 3.9% of the labeled nucleotides of the training slice lie in a peak, so we give the positive nucleotides the larger weight $w_+ = 49.06$, which we chose together with the other hyperparameters (Section 2.9). Both kinds of negative examples enter $\mathcal{L}_{\text{nt}}$, and all of their nucleotides have the label 0. The pair term is the binary cross-entropy of the pair logit,

$$\mathcal{L}_{\text{pair}} = -\frac{1}{B} \sum_{i=1}^{B} \Big[ c_i \log \sigma(g_i) + (1 - c_i) \log\big(1 - \sigma(g_i)\big) \Big], \tag{S8}$$

where $c_i$ is the label of the example and $B$ is the number of examples in the batch. The pair logit is the output that we evaluate. The nucleotide term gives the head a dense training signal at the positions of the peaks, which the pair label alone does not give.

We train with the nucleotide term alone for the first $T_0 = 15{,}400$ steps, so that the nucleotide logits learn where the peaks are before the pair term is added. Then $\lambda$ grows linearly to $\lambda_{\max} = 10$ over the next $T_1 = 15{,}400$ steps and stays at this value until the end of training,

$$\lambda(t) = \lambda_{\max} \min\Big(1, \max\Big(0, \frac{t - T_0}{T_1}\Big)\Big). \tag{S9}$$

While $\lambda = 0$, the scalars $w$ and $b$ get no gradient, but the scale of the nucleotide logits changes. At step $T_0$, we therefore set $w$ and $b$ once from 2,048 training examples that we score with dropout turned off. We set $w = 1/\text{sd}(z)$ and $b = \text{logit}(\pi) - w\,\bar{z}$. Here $\bar{z}$ and $\text{sd}(z)$ are the mean and the standard deviation of the pooled scores of these examples, and $\pi = 0.481$ is the share of positive examples in the training slice. When the pair term is added, the pair logits therefore have unit standard deviation, and their mean is the logit of the positive rate.

#### 2.7 Batch sampling and optimization

We sample the examples of each batch so that the proteins and the labels are balanced. Two properties of the training slice make this necessary. First, the number of training examples per protein, $n_p$, ranges from 99 to 32,238. Uniform draws would let a few proteins dominate the gradient, and an equal number of draws for each protein would repeat the examples of the rarest proteins many times. We therefore give protein $p$ a number of draws proportional to $n_p^{1/2}$. Second, the share of positive examples differs between proteins, so the pair logit could learn how often a protein binds instead of whether the given RNA binds it. We therefore draw half of the examples of each protein from its positives and half from its negatives. We draw without replacement and train for 154,000 steps, so every training example is drawn at least once.

A batch holds at most 65,536 tokens and 32 examples. To keep padding low, we order the drawn examples by length in buckets of 64 tokens, in random order within each bucket. We then cut this list into batches and shuffle the batches. We do not sort the examples fully by length. The length of an input depends on the length of the protein, so a full sort would put the examples of one protein into the same batch.

We use AdamW [17] with a learning rate of $5 \times 10^{-4}$ and a weight decay of 0.01. The learning rate grows linearly over the first 5% of the steps and then follows a cosine decay to zero. We clip the norm of the gradient at 1.0. Training takes 12 hours on one NVIDIA RTX PRO 6000 GPU. We chose the hyperparameters in short training runs that we scored on the validation slices (Section 2.9).

**Table S4.** Hyperparameter search. Each run has 3,000 steps. Score: mean protein-macro AUROC over the three validation slices. Each step starts from the values kept in the earlier steps. Bold: kept value.

| Value | Score | Value | Score |
|---|---|---|---|
| *1. Learning rate* | | *4. Exponent $e$ of the protein draws* | |
| $2 \times 10^{-4}$ | 0.597 | $e = -1$ | 0.610 |
| $\mathbf{5 \times 10^{-4}}$ | **0.604** | $e = -1/2$ | 0.635 |
| $1.5 \times 10^{-3}$ | 0.597 | $e = 0$ | 0.636 |
| *2. $\lambda_{\max}$ and $w_+$* | | $\mathbf{e = 1/2}$ | **0.661** |
| $\lambda_{\max} = 0.2$, $w_+ = 49.06$ | 0.604 | *5. Pair label* | |
| $\lambda_{\max} = 0.2$, $w_+ = 7$ | 0.617 | **Hard** | **0.661** |
| $\lambda_{\max} = 0.6$, $w_+ = 49.06$ | 0.617 | Graded | 0.653 |
| $\lambda_{\max} = 0.6$, $w_+ = 7$ | 0.626 | *6. $\lambda_{\max}$, mean of two seeds* | |
| $\boldsymbol{\lambda_{\max} = 2}$, $\mathbf{w_+ = 49.06}$ | **0.635** | $\lambda_{\max} = 2$ | 0.644 |
| $\lambda_{\max} = 2$, $w_+ = 7$ | 0.636 | $\lambda_{\max} = 5$ | 0.656 |
| *3. $T_0$, share of the steps* | | $\boldsymbol{\lambda_{\max} = 10}$ | **0.665** |
| 3% | 0.629 | | |
| **10%** | **0.635** | | |
| 25% | 0.630 | | |

#### 2.8 Model selection and decision threshold

Every 10,000 steps, we score a fixed random sample of 10,000 examples from each validation slice, with dropout turned off. For each slice, we compute the protein-macro AUROC. For each protein that has at least 10 positive and 10 negative examples, we compute the AUROC of the model on the examples of this protein, and we take the unweighted mean over these proteins. We keep the checkpoint with the highest mean protein-macro AUROC over the three validation slices, so each slice counts the same regardless of its size. This is the checkpoint of step 154,000.

We then score all examples of the three validation slices with the selected checkpoint. We choose the decision threshold that maximizes the Matthews correlation coefficient (MCC), averaged over the three slices with equal weight. Every predicted probability on the validation slices is a candidate threshold. The chosen threshold is 0.439. Only after this step do we score the three test slices, once and on all of their examples.

#### 2.9 Hyperparameter search

We chose six hyperparameters in short training runs. They are the learning rate, the final weight $\lambda_{\max}$ of $\mathcal{L}_{\text{pair}}$, the weight $w_+$ of positive nucleotides, the length $T_0$ of the phase with $\mathcal{L}_{\text{nt}}$ alone, the exponent $e$ of the protein draws, and the form of the pair label. Each run has 3,000 steps. The phase with $\mathcal{L}_{\text{nt}}$ alone and the ramp of $\lambda$ take 10% of the steps each, as in the full schedule. The learning rate grows over the first 5% of the steps and decays to zero at step 3,000, and the examples are drawn with replacement. At the end of each run, we score a fixed random sample of 10,000 examples from each validation slice. The score of a run is the mean protein-macro AUROC over the three validation slices (Section 2.8). The test slices are not scored in the search.

We search the hyperparameters in six steps, in the order of Table S4. Each step starts from the values that the earlier steps kept. The starting values are a learning rate of $5 \times 10^{-4}$, $\lambda_{\max} = 0.2$, $w_+ = 49.06$, $T_0$ of 10% of the steps, $e = -1/2$, and the hard pair label. The starting value of $w_+$ is the ratio of labeled nucleotides outside and inside peaks in the batches of the starting setting. To measure the noise of the score, we trained one setting with three random seeds. The standard deviation of its score over the seeds is 0.005. We keep a new value only when its score is higher than that of the current value by more than 0.015, which is twice the standard deviation of the difference between two runs. When each value has two runs, this margin is 0.011.

Table S4 gives the scores. Learning rates of $2 \times 10^{-4}$ and $1.5 \times 10^{-3}$ gave lower scores than $5 \times 10^{-4}$. The weight $\lambda_{\max}$ had the largest effect. With $w_+ = 49.06$, the score rose from 0.604 at $\lambda_{\max} = 0.2$ to 0.635 at $\lambda_{\max} = 2$. A weight of $w_+ = 7$ scored 0.013, 0.009, and 0.002 higher at $\lambda_{\max} = 0.2$, 0.6, and 2. This difference became smaller as $\lambda_{\max}$ grew, and it was below the margin at every $\lambda_{\max}$, so we kept $w_+ = 49.06$. A $T_0$ of 3% or 25% of the steps lowered the score. For the protein draws, $e = 1/2$ gave the highest score, 0.027 above the starting value $e = -1/2$. The graded pair label gives a positive example a target that grows from 0.6 to 1 with the number of its peak nucleotides and reaches 1 at 220 nt. It scored lower than the hard label. The score still rose at $\lambda_{\max} = 2$, so in the last step we trained $\lambda_{\max} = 2$, 5, and 10 with two seeds each. The mean scores were 0.644, 0.656, and 0.665, and we kept $\lambda_{\max} = 10$.

### 3 Compared methods

This section gives the details of the compared methods in Section 2.4.1 of the main text. Sections 3.1 to 3.5 describe the five methods that we retrain, and Section 3.6 explains why we did not retrain four other methods.

We train each of the five methods on the training slice, with the same examples and the same negatives as RPI-Zero. We keep the features, the architecture, and the training procedure of each method as its paper and its released code describe them. Where a method trains for a fixed number of epochs or selects its model on cross-validation folds, we instead select it on the three validation slices or stop its training early on them, as we do for RPI-Zero. We then score each method once on all examples of the three test slices. Each subsection ends with the decision threshold of the method, which the threshold-based metrics in Table 2 of the main text use.

#### 3.1 IPMiner

IPMiner [20] uses k-mer frequencies, as many earlier methods do, and these frequencies can be computed for an RNA of any length. It describes the RNA by its 4-mer frequencies and the protein by its conjoint triad frequencies [23], which count amino acid triplets over a reduced alphabet of seven groups. Both are computed on the full sequences, so no part of an example is cut. A stacked autoencoder with three layers learns a code for each molecule, and the two encoders are then fine-tuned together with a classification layer. Three random forests use the fine-tuned codes, the autoencoder codes, and the raw frequencies, and a logistic regression combines their outputs.

The paper does not say how the training data of the logistic regression are made. We train it on out-of-fold predictions of a 3-fold cross-validation on the training slice. In each fold, we fit the autoencoders, the fine-tuned network, and the random forests again, so that no out-of-fold prediction comes from a model that saw the example in training. The released code fits the feature scaling on the training and test examples together, and we fit it on the training slice only. Where the paper and the released code differ, we follow the paper, for example in the tied weights of the encoder and the decoder. We fine-tune for 100 epochs and keep the epoch with the lowest cross-entropy averaged over the three validation slices. The decision threshold is 0.5.

#### 3.2 NPI-GNN

We include NPI-GNN [24] to test a method that uses a graph of known interactions on molecules that have no interaction in the training data. NPI-GNN builds a graph whose nodes are the RNAs and the proteins. As in the released code, the edges are all training pairs, both positive and negative, and the RNA of each example is its own node. For each pair, NPI-GNN takes the subgraph made of the two nodes and their direct neighbors. A node is described by its role in the subgraph, by a 64-dimensional node2vec embedding [8] of the training graph, and by k-mer frequencies. These are the 3-mer frequencies of the RNA and the 2-mer frequencies of the protein over a reduced alphabet. Three GraphSAGE layers [9], each followed by top-$k$ pooling, classify the subgraph.

As in the released code, a molecule that has no edge in the training graph gets a zero node2vec vector and no neighbors. Every test pair has at least one such molecule. If neither molecule of a pair is in the training graph, the model gives the pair a probability of about 0.36 and predicts it as negative. This is the case for all examples of the unseen pair slice, 73% of the examples of the unseen protein slice, and 15% of the examples of the unseen RNA slice. We train for 50 epochs with Adam, as in the released code, and save a checkpoint every 5 epochs. We keep the checkpoint with the highest MCC at a threshold of 0.5, averaged over the three validation slices. The released code selects the epoch on the test folds of a cross-validation, which we avoid. The decision threshold is 0.5.

#### 3.3 ZHMolGraph

The authors of ZHMolGraph [15] tested it on RNAs and proteins that are absent from training. ZHMolGraph embeds the RNA with RNA-FM [4] and the protein with ProtT5 [7], and it takes the mean over the tokens of each sequence. RNA-FM reads at most 1,022 nt, so a longer RNA is cut into segments, and the segment embeddings are averaged as in the released code. A GraphSAGE network [9] is trained without labels on a graph of the RNAs and the proteins. As in the released code, its edges are the pairs, positive and negative, whose RNA and protein both appear in positive training pairs, so no test pair is an edge. The node embeddings of this network are joined with the language-model embeddings and passed to a feed-forward classifier. We train the classifier with Adam, a constant learning rate, and a batch size of 64, as in the released code. Every 10,000 steps, we score fixed samples of about 10,000 examples from each validation slice. The score is the binary cross-entropy averaged first over the proteins of a slice and then over the three slices. We stop after five checks without improvement. The decision threshold is 0.392, which maximizes the MCC on the validation slices.

#### 3.4 CORAL

The authors of CORAL [3] tested it on splits in which no test RNA or protein is similar to the training data, which is the setting of our benchmark. CORAL encodes the RNA with DNABERT-2 [35] and the protein with the ESM-2 model of 150 million parameters [14], and it joins the two with two cross-attention layers. DNABERT-2 has no token for U, so we write each U as T, as the authors do for their own data.

We train with the settings of the paper. The loss is the sum of the classification loss and a masked language modeling loss on 15% of the tokens of both sequences. Both encoders are fine-tuned with LoRA of rank 8, and the cross-attention layers and the output heads are fully trained. We use AdamW with a learning rate of $10^{-6}$, a batch size of 8, and a linear schedule with a warmup of 5% over a budget of 10 epochs. The data of CORAL have no protein longer than 1,250 residues, and its inference code cuts longer proteins to 1,250 tokens. We therefore drop the training examples with longer proteins, which are 9.1% of the training slice, and cut longer proteins to 1,250 tokens in the validation and test slices. Every 10,000 steps, we compute the cross-entropy on fixed samples of 10,000 examples from each validation slice, and we stop after five checks without improvement. The decision threshold is 0.5.

#### 3.5 ZHMolTopoRPI

The authors of ZHMolTopoRPI [5] also tested it on RNAs and proteins that are absent from training. ZHMolTopoRPI describes the RNA by topological invariants of the positions of each 4-mer along the sequence, which it computes with persistent Stanley–Reisner theory. A variance filter and a principal component analysis reduce these 9,984 features to 3,240, and we fit both on the training slice only. The protein is the mean of the last layer of the ESM-2 model of 3 billion parameters [14] over all residues of the full-length protein. A network with one tower for each molecule and a gated fusion of the two towers gives the prediction. It is trained with a focal loss [13] and a supervised contrastive loss [12].

Where the paper and the released code differ, we follow the paper. We do not scale the features after the principal component analysis, and the gate has one value for each feature instead of a single value. The released code also cuts proteins at 1,022 residues before ESM-2, while we use the full protein. The paper reports hyperparameters for each of its data sets, and we use the values of its NPInter2 data set. We stop training when the validation loss has not improved for five epochs. We keep the epoch with the highest MCC at a threshold of 0.5, averaged over the three validation slices. As in the paper, the decision threshold maximizes the MCC. We choose it for each test slice on the matching validation slice, which gives 0.50 for unseen RNAs and 0.49 for unseen proteins and unseen pairs. The released code chooses the threshold on the evaluation fold, which we avoid.

#### 3.6 Methods that we did not retrain

We examined four more methods that predict RNA-protein interaction from the two sequences. We did not retrain them, for the reasons below.

**3.6.1 BioLLMNet, RPIPLM, and PaRPI:** These methods read only short RNAs. BioLLMNet [1] describes the RNA by one embedding from RiNALMo [22], which reads at most 1,022 nt. RPIPLM [16] encodes the RNA with RNABERT [2], which has 440 position embeddings and accepts only sequences shorter than 440 nt. PaRPI [21] reads a window of 101 nt in training and in testing, and this length is fixed in its code. The RNA of a training example has a median length of 2,711 nt. Of the 1,169,793 training examples, only 171,191 (14.6%) have an RNA of at most 1,022 nt, 44,454 (3.8%) have an RNA shorter than 440 nt, and 17,855 (1.5%) have an RNA of at most 101 nt. None of the three papers describes how to score a longer RNA, for example by cutting it into windows and combining the scores of the windows. A windowing step of our own would add choices that the authors did not make, so its results would not be the results of the published method. ZHMolGraph has the same limit of 1,022 nt through RNA-FM, but its released code cuts a long RNA into segments, so we could keep its own procedure (Section 3.3).

**3.6.2 The attention model of Yu et al.:** Yu et al. [32] released three trained models, one for each of their protein encodings, and code for prediction only. The training code is not public, and the released models do not match the paper. They have 4 attention heads instead of 8 and a different scale of the attention scores. They also differ from the paper in the number of filters of one convolution, in the dropout, in the learning rate schedule, and in the padding of the ProteinBERT features. The structure-aware model uses a one-dimensional convolution where the paper describes a three-dimensional one. A model that follows the paper would therefore be a model that the authors never trained.

### 4 Additional experiments and results

This section adds two analyses to Section 3 of the main text. Section 4.1 compares the ranks of the six methods across the three test slices. Section 4.2 measures what three parts of RPI-Zero add to its predictions.

#### 4.1 Ranks of the methods on the test slices

Fig. S1 gives the rank of each method on the eight metrics in Table 2 of the main text, for each of the three test slices. RPI-Zero ranks first in all 24 rows. The order of the other methods changes from slice to slice. By AUROC, ZHMolTopoRPI is second on unseen RNAs but fourth on unseen proteins. IPMiner is second on unseen proteins but fourth on unseen pairs, and CORAL is second on unseen pairs but fourth on unseen RNAs. Over the 24 rows, the mean rank is 1.0 for RPI-Zero, 2.6 for ZHMolTopoRPI, 3.3 for IPMiner, and 3.5 for CORAL.

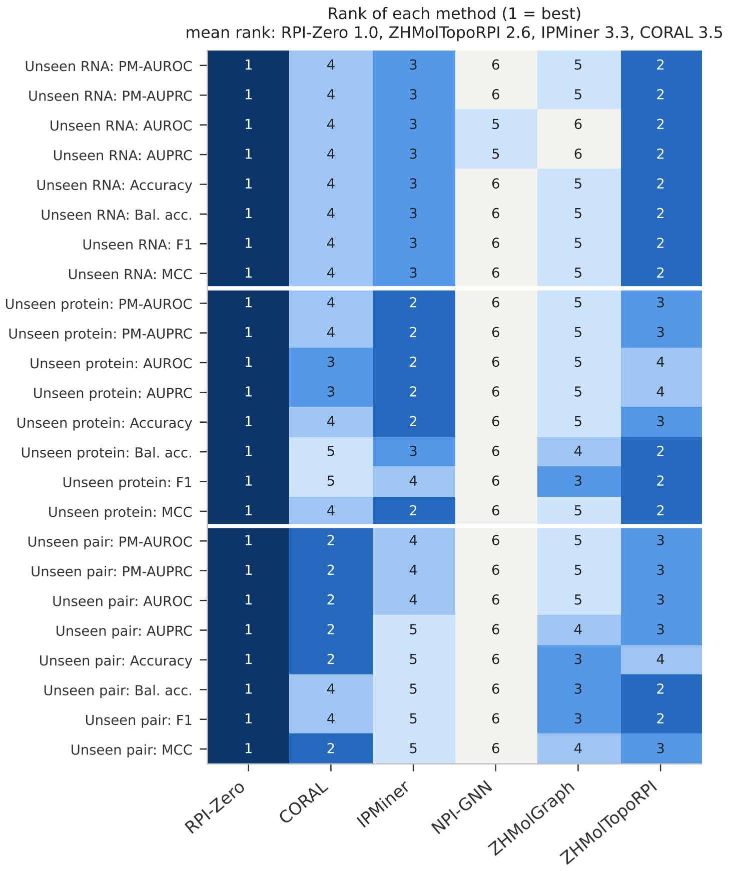


**Fig. S1.** Rank of each method on the three test slices (1 = best). Each row is one metric on one slice, with the values in Table 2 of the main text. Tied methods share the better rank. The title gives the mean rank over the 24 rows for the four best methods. PM: protein-macro. Bal. acc.: balanced accuracy.

### 4.2 Contribution of the architecture and the losses

We test three parts of RPI-Zero, which are the convolution blocks, the nucleotide loss $\mathcal{L}_{\text{nt}}$, and the pair loss $\mathcal{L}_{\text{pair}}$. For each part, we train a model without it and compare this model with the full model.

**4.2.1 Ablated models:** Without the convolution blocks, the head has none of the five blocks of Eq. (S2), so $\mathbf{h}_j = \mathbf{h}_j^{(0)}$. The logit of a nucleotide then depends only on the RNA-X embedding of the same nucleotide, and the protein context $\mathbf{q}$ no longer gates the RNA features. The motif score, the protein term, the RNA term, and the pooling stay as in the full model. Without $\mathcal{L}_{\text{nt}}$, the loss is $\lambda(t)\,\mathcal{L}_{\text{pair}}$, and the labels of the nucleotides are not used. There is no phase with $\mathcal{L}_{\text{nt}}$ alone, and $\lambda$ grows from 0 to 10 over the first 3,000 steps. The head still computes the nucleotide logits, but they are trained only through the pooling. Without $\mathcal{L}_{\text{pair}}$, the loss is $\mathcal{L}_{\text{nt}}$ alone, and the labels of the examples are not used. The pair logit is still computed from the nucleotide logits by the pooling of Eq. (S5).

**4.2.2 Training and evaluation:** Each ablation needs its own training run, so all runs in this section use a shorter schedule of 30,000 steps. The phase with $\mathcal{L}_{\text{nt}}$ alone and the ramp of $\lambda$ take 3,000 steps each, and the examples are drawn with replacement. As the control, we train the full model again with this schedule. All other settings are those of Table S3, and all runs use the same random seed. Every 2,000 steps, we score the fixed samples of the validation slices (Section 2.8) and keep the checkpoint with the highest mean protein-macro AUROC over the three slices. We then compute the protein-macro AUROC of this checkpoint on all examples of each validation slice.

We measure localization with the within-example AUROC. For a positive example, this is the AUROC of its nucleotide logits against the labels of its nucleotides. It compares only nucleotides of the same RNA, so a score that does not localize the peaks gives 0.5. We average it over the positive examples of each protein and then over the proteins. On the validation slices, we compute it on the positive examples of the fixed samples. For each difference from the control, we compute a 95% confidence interval from 10,000 bootstrap resamples of the proteins. We test the differences with a two-sided Wilcoxon signed-rank test [29] and correct the $p$-values with the Holm method [11], as in Section 3.1 of the main text. We call a difference significant when its interval excludes zero and its corrected $p$-value is below 0.05.

**Table S5.** Ablations of RPI-Zero on the validation slices. All models are trained for 30,000 steps. The first row is the full model, and each other row removes one part of it. The protein-macro AUROC of the pair logit is computed on all examples of a slice, and the difference from the full model is given in parentheses. All nine differences are significant, with a 95% bootstrap confidence interval below zero and a Holm-corrected Wilcoxon $p$-value below 0.05. The within-example AUROC ranks the nucleotides of each positive example by their logits, and a random score gives 0.5. RNA, Protein, and Pair are the unseen RNA, unseen protein, and unseen pair slices.

| | Protein-macro AUROC | | | Within-example AUROC | | |
|---|---|---|---|---|---|---|
| Model | RNA | Protein | Pair | RNA | Protein | Pair |
| RPI-Zero | 0.751 | 0.698 | 0.691 | 0.724 | 0.663 | 0.644 |
| Without convolution blocks | 0.721 (−0.030) | 0.654 (−0.044) | 0.655 (−0.035) | 0.653 | 0.600 | 0.593 |
| Without $\mathcal{L}_{\text{nt}}$ | 0.730 (−0.022) | 0.674 (−0.023) | 0.673 (−0.018) | 0.552 | 0.525 | 0.523 |
| Without $\mathcal{L}_{\text{pair}}$ | 0.717 (−0.034) | 0.650 (−0.048) | 0.649 (−0.042) | 0.764 | 0.670 | 0.651 |

##### 4.2.3 Results on the validation slices:

Table S5 gives the results. Removing any of the three parts lowers the protein-macro AUROC on all three validation slices, and all nine drops are significant. The drop is 0.030 to 0.044 without the convolution blocks, 0.018 to 0.023 without $\mathcal{L}_{\text{nt}}$, and 0.034 to 0.048 without $\mathcal{L}_{\text{pair}}$. For each part, the largest drop is on unseen proteins. On this slice, the models without the convolution blocks, without $\mathcal{L}_{\text{nt}}$, and without $\mathcal{L}_{\text{pair}}$ are below the full model for 22, 18, and 21 of the 23 proteins.

The convolution blocks and $\mathcal{L}_{\text{nt}}$ are also needed to localize the binding sites. Without $\mathcal{L}_{\text{nt}}$, the within-example AUROC falls from 0.724, 0.663, and 0.644 to 0.552, 0.525, and 0.523, which is close to the 0.5 of a random score. The label of the example alone therefore does not teach the head where the protein binds. Without the convolution blocks, the within-example AUROC falls to 0.653, 0.600, and 0.593. Without $\mathcal{L}_{\text{pair}}$, $\mathcal{L}_{\text{nt}}$ still trains the nucleotide logits, and the within-example AUROC does not fall. The protein-macro AUROC still falls by 0.034 to 0.048 in this case. Good localization alone therefore does not give a good pair score, and $\mathcal{L}_{\text{pair}}$ trains the head to turn the nucleotide logits of a whole RNA into one score for the pair.

##### 4.2.4 Results on the test slices:

We also scored the three test slices once with the selected checkpoint of each run. Without the convolution blocks, the protein-macro AUROC on unseen RNAs, unseen proteins, and unseen pairs falls by 0.034, 0.023, and 0.023. Without $\mathcal{L}_{\text{pair}}$, it falls by 0.037, 0.036, and 0.031. All six drops are significant. We also compute the within-example AUROC on all positive examples of the test slices. Without $\mathcal{L}_{\text{nt}}$, it falls from 0.694, 0.666, and 0.675 to 0.541, 0.531, and 0.529. It is lower than that of the full model for all 184 proteins of the unseen RNA slice, all 24 proteins of the unseen protein slice, and 23 of the 24 proteins of the unseen pair slice. Without the convolution blocks, it falls by 0.066 to 0.071 on the three test slices.

#### 4.3 Similarity between training and test data in the published evaluations

The authors of the five compared methods evaluated them on other data sets, and most of these papers report much higher scores than those in Table 2 of the main text. We therefore measured how many test RNAs and test proteins of these evaluations are unseen under the rules of our benchmark. Here, an evaluation is one way in which a paper divides its data into training and test sets. Examples are a 5-fold cross-validation on one data set, or training on one data set and testing on another. For each evaluation, we compared its original test set with its original training set. We took both sets from the released split files, or we rebuilt them by running the released code that makes the folds. The rebuilt data sets have the pair counts reported in the papers, except for one or two duplicate pairs that the released code itself merges. Table S6 gives the results for 24 evaluations. These are 16 cross-validations, four splits of CORAL, and four tests on another data set.

**4.3.1 Seen and unseen molecules:** For each evaluation, we cluster all distinct RNA sequences and all distinct protein sequences, from the training and the test set together, with the MMseqs2 [26] settings of our benchmark (Section 1.4). On the transcripts and proteins of our benchmark, these settings give the same 26,684 RNA clusters and 181 protein clusters. A test RNA is *seen* if its cluster holds an RNA of a training pair of the same fold, and it is *unseen* otherwise. We count the RNAs of both positive and negative training pairs, because a model reads both. Proteins are treated in the same way. A test molecule with an identical sequence in training is therefore always seen.

We then put each test pair into one of four groups. An unseen RNA pair has an unseen RNA and a seen protein, as in the unseen RNA slice of our benchmark. An unseen protein pair has a seen RNA and an unseen protein, as in the unseen protein slice. An unseen pair has an unseen RNA and an unseen protein, as in the unseen pair slice. A both-seen pair has a seen RNA and a seen protein, and our benchmark has no test slice for such pairs. Our benchmark keeps each cluster in one bucket. In its unseen RNA and unseen pair slices, every RNA is therefore unseen in this sense. In its unseen protein and unseen pair slices, every protein is unseen. A test pair that does not occur in the training data is not an unseen pair unless both of its molecules are unseen.

A 5-fold cross-validation has five folds, and each fold has its own training set and test set. We compare the test set of each fold with the training set of the same fold. We then add the counts of the five folds and divide them by the total number of test molecules or test pairs of the five folds. We do the same for repeated cross-validations. CORAL released five versions of each of its four splits, and we add their counts in the same way.

**4.3.2 Cross-validation:** Cross-validation is the main evaluation of IPMiner, NPI-GNN, ZHMolGraph, and ZHMolTopoRPI. It divides the pairs of a data set into folds without any control of sequence similarity, so the same RNA or protein can occur in a training fold and in a test fold. In the 16 cross-validations of Table S6, 82.2% to 99.9% of the test pairs are both-seen pairs, with a median of 95.9%. No cross-validation has more than 7.4% unseen pairs. NPInter2 and RPI7317 are the main data sets of NPI-GNN, ZHMolGraph, and ZHMolTopoRPI. IPMiner and NPI-GNN use the name NPInter2.0 for NPInter2. On these two data sets, no test protein is unseen, and at most 4.0% of the test RNAs are unseen. On NPInter2, the three methods report MCCs of 0.868, 0.911, and 0.923 [5, 15, 24].

IPMiner also reports a cross-validation on NPInter2.0. We do not include it in Table S6, because the proteins of its negative pairs were drawn at random and cannot be recovered. Each of these negative pairs uses the RNA of a positive pair, and the released code places the two pairs in different folds. In every fold, every test RNA is therefore also the RNA of a training pair.

**4.3.3 Tests on molecules from another data set:** IPMiner trains a model on RPI488. RPI488 has only 25 distinct RNA sequences. IPMiner tests this model on NPInter2.0 and on RPIntDB. Most test pairs of these two tests are unseen pairs, 98.8% and 65.6%. Both test sets hold only binding pairs, and IPMiner reports the share of them that it predicts as binding, which is 96.7% and 86.4% [20]. A model that predicts every pair as binding would reach 100% on both.

**Table S6.** Unseen test molecules and groups of test pairs in the published evaluations of the five compared methods. Each test set is compared with the training set of the same evaluation, with the clustering rules of our benchmark. A test molecule is unseen if no molecule of its cluster occurs in a training pair. Unseen RNA pairs: unseen RNA and seen protein. Unseen protein pairs: seen RNA and unseen protein. Unseen pairs: unseen RNA and unseen protein. Both-seen pairs: seen RNA and seen protein. The four groups of pairs add up to 100%. CV: 5-fold cross-validation, with repeated runs pooled. The thresholds of the CORAL splits are the identities for RNA and protein. *The test set holds only binding pairs.

| | | Unseen test molecules (%) | | Test pairs (%) | | | |
|---|---|---|---|---|---|---|---|
| Method | Evaluation | RNAs | Proteins | Unseen RNA pairs | Unseen protein pairs | Unseen pairs | Both-seen pairs |
| CORAL | Pairwise split, 80%/60% | 28.6 | 27.5 | 18.4 | 12.1 | 4.3 | 65.2 |
| | Pairwise split, 50%/30% | 29.2 | 28.0 | 18.5 | 12.0 | 4.4 | 65.1 |
| | Component-wise split, 80%/60% | 95.2 | 78.4 | 18.4 | 4.9 | 75.0 | 1.7 |
| | Component-wise split, 50%/30% | 94.9 | 97.7 | 1.6 | 5.8 | 92.5 | 0.1 |
| IPMiner | CV, RPI488 | 1.9 | 16.4 | 0.2 | 16.7 | 0.2 | 82.9 |
| | CV, RPI369 | 13.7 | 19.8 | 3.0 | 8.5 | 6.1 | 82.4 |
| | CV, RPI1807 | 9.1 | 9.5 | 0.9 | 1.6 | 1.6 | 95.9 |
| | Trained on RPI488, tested on NPInter2.0* | 100.0 | 97.9 | 1.2 | 0.0 | 98.8 | 0.0 |
| | Trained on RPI488, tested on RPIntDB* | 98.8 | 58.4 | 7.9 | 2.3 | 65.6 | 24.2 |
| NPI-GNN | CV, NPInter2.0 | 3.8 | 0.0 | 3.5 | 0.0 | 0.0 | 96.5 |
| | CV, RPI7317 | 0.3 | 0.0 | 0.3 | 0.0 | 0.0 | 99.7 |
| | CV, RPI2241 | 6.2 | 6.1 | 0.8 | 2.2 | 1.1 | 95.9 |
| | CV, RPI369 | 13.7 | 20.9 | 2.6 | 8.9 | 6.1 | 82.4 |
| | CV, NPInter2.0 lncRNA | 0.1 | 0.0 | 0.1 | 0.0 | 0.0 | 99.9 |
| ZHMolGraph | CV, NPInter2 | 4.0 | 0.0 | 3.7 | 0.0 | 0.0 | 96.3 |
| | CV, RPI7317 | 0.4 | 0.0 | 0.4 | 0.0 | 0.0 | 99.6 |
| | Trained on NPInter2, tested on TheNovel | 52.3 | 92.3 | 11.8 | 28.5 | 46.5 | 13.2 |
| ZHMolTopoRPI | CV, NPInter2 | 3.9 | 0.0 | 3.6 | 0.0 | 0.0 | 96.4 |
| | CV, RPI7317 | 0.3 | 0.0 | 0.3 | 0.0 | 0.0 | 99.7 |
| | CV, RPI488 | 1.8 | 18.6 | 0.0 | 17.2 | 0.6 | 82.2 |
| | CV, RPI1807 | 19.0 | 21.1 | 4.0 | 5.2 | 7.4 | 83.4 |
| | CV, RPI2241 | 6.4 | 9.5 | 2.4 | 5.7 | 1.6 | 90.3 |
| | CV, NPInter v2.0 | 4.2 | 10.5 | 3.6 | 2.1 | 0.0 | 94.3 |
| | Trained on NPInter2, tested on TheNovel* | 52.3 | 92.3 | 19.9 | 11.1 | 46.7 | 22.3 |

ZHMolGraph and ZHMolTopoRPI test on TheNovel. Their authors describe it as a set of unknown nodes, whose RNAs and proteins are absent from training [5, 15]. TheNovel holds the interactions of NPInter5 [34] whose RNA and protein names do not occur in NPInter2. It is scored by the five models of the NPInter2 cross-validation, and we compare each model with its own training data. No sequence of TheNovel is identical to a training sequence. Under the rules of our benchmark, however, 47.7% of its RNAs and 7.7% of its proteins are seen, and only 46.5% of its pairs are unseen pairs. The other pairs are unseen protein pairs (28.5%), unseen RNA pairs (11.8%), and both-seen pairs (13.2%). The positive pairs that ZHMolTopoRPI released give almost the same share of unseen pairs, 46.7%.

**4.3.4 CORAL:** Of the five papers, CORAL has the evaluation that is closest to our benchmark. Its component-wise splits assign RNA clusters and protein clusters to training or test, and they drop every pair that has one molecule on each side [3]. At thresholds of 50% for RNA and 30% for protein, 92.5% of the test pairs are unseen pairs under our rules. CORAL reports its main results at 80% for RNA and 60% for protein [3]. With these thresholds, 21.6% of the test proteins are seen at the 30% identity of our benchmark, and 75.0% of the test pairs are unseen pairs. Because the mixed pairs are dropped, the component-wise splits are not designed to test an unseen RNA with a seen protein, or a seen RNA with an unseen protein. Our benchmark tests these two cases in its unseen RNA and unseen protein slices. The pairwise splits of CORAL keep the mixed pairs, but in these splits 65% of the test pairs are both-seen pairs and only 4.3% to 4.4% are unseen pairs.

**4.3.5 Check with a direct comparison:** Two similar sequences can fall into different clusters. We therefore also compared each test sequence directly with all training sequences of the same evaluation, with the same thresholds. This check can only turn unseen molecules into seen ones. Except on TheNovel, it lowers the shares of unseen molecules by less than 3 percentage points. On TheNovel, the share of unseen RNAs falls from 52.3% to 41.9%.

#### 4.4 Stratified evaluation by RNA and protein type and length

Table 2 of the main text gives one value per test slice. Here, we test whether the lead of RPI-Zero holds for different kinds of RNAs and proteins. The test transcripts range from 47 nt to more than 100,000 nt, and the test proteins from 127 to 2,804 amino acids. Many earlier methods were built for one type of RNA (Section 1 of the main text). We therefore divide the test examples into groups by the type and the length of the RNA and by the class and the length of the protein, and we compare the six methods within each group.

We use the three test slices together, because the unseen protein and unseen pair slices hold only 24 proteins. If we divide these 24 proteins by class or by length, 7 of the 8 groups would hold fewer than 10 proteins. The 184 proteins of the unseen RNA slice are scored on their unseen RNA examples, and the 24 test proteins are scored on their unseen protein and unseen pair examples.

The type of an RNA is the GENCODE biotype of its transcript [19]. We report protein-coding RNAs and lncRNAs, which together hold 94% of the test examples. The length of an RNA is the length of its whole spliced transcript. We do not use the length of the RNA in the example. A transcript that is longer than the budget of the protein is cut into parts (Section 1.3), so the length of a part depends on the protein. The length of a protein is its number of amino acids. One border of the protein length groups lies at 1,250 amino acids, which is the length limit of CORAL (Section 3.4).

The class of a protein describes what decides where the protein binds. We grouped the proteins by hand into four classes, and Table S7 lists the proteins of each class. A motif-driven protein binds a short sequence motif in the RNA, as PTBP1 and RBFOX2 do. For a non-sequence-driven protein, the binding site is decided mainly by something other than the RNA sequence. This class covers three cases. First, some proteins add, remove, or read a chemical modification of the RNA, such as m6A. In the input sequence, a modified nucleotide looks the same as an unmodified one, so the input does not show the modification. Second, some proteins belong to the miRNA pathway. AGO2, for example, is guided to its targets by a miRNA, and this miRNA is not part of the input. Third, some proteins bind RNA as part of a large complex, such as the spliceosome, and the complex decides where they bind. A fixed-target protein binds a few specific RNAs, such as rRNAs or snRNAs. The fourth class holds all other proteins, whose mechanism is mixed or unclear.

The metric is the protein-macro AUROC. In each group, we compute the AUROC of a method on the examples of every protein that has at least 10 positive and 10 negative examples in the group, and we take the mean over these proteins. We use the AUROC because its random baseline is 0.5 in every group, while that of the AUPRC depends on the share of positives. As in Section 3.1 of the main text, but with the AUROC, we compare RPI-Zero with each method. We use 10,000 bootstrap resamples of the proteins and a two-sided Wilcoxon signed-rank test [29]. We correct the $p$-values with the Holm method [11], separately for each property. Table S8 gives the results.

RPI-Zero has the highest protein-macro AUROC in all 15 groups. Its lead over the second-best method of a group ranges from 0.011 to 0.051. In 14 of the 15 groups, the difference from each of the five methods has a 95% confidence interval above zero and a corrected $p$-value below 0.05. The exception is lncRNAs, where the lead over IPMiner, CORAL, and ZHMolTopoRPI is not significant.

IPMiner and NPI-GNN were built for interactions of proteins with ncRNAs [20, 24]. On lncRNAs, however, they reach 0.645 and 0.494, against 0.656 for RPI-Zero. The gap is larger for unseen proteins. On the lncRNAs of the unseen protein and unseen pair slices, RPI-Zero reaches 0.707 and 0.654, and the best other method reaches 0.668 (IPMiner) and 0.613 (CORAL).

RPI-Zero is the best method in all four protein classes. This includes the non-sequence-driven proteins, whose binding sites are not decided mainly by the RNA sequence. On these proteins, RPI-Zero reaches 0.726, and the second-best method, IPMiner, reaches 0.698.

Protein length shows a weakness of CORAL. Its protein-macro AUROC drops from 0.662 and 0.682 for proteins shorter than 800 amino acids to 0.622 and 0.620 for longer proteins. Proteins longer than 1,250 amino acids are also cut by CORAL, which was trained only on shorter proteins. RPI-Zero reads the whole protein, and its protein-macro AUROC stays between 0.730 and 0.767 in all four length groups. On proteins longer than 1,250 amino acids, RPI-Zero is 0.110 above CORAL.

**Table S7.** Classes of the 208 proteins of the test slices. Of these proteins, 204 have at least 10 positive and 10 negative test examples and enter Table S8.

| Class | $n$ | Proteins |
|---|---|---|
| Motif-driven | 48 | CELF2, CNBP, CPEB4, CPSF6, CSTF2, CSTF2T, ELAVL1, FUBP3, HNRNPA1, HNRNPA2B1, HNRNPD, HNRNPF, HNRNPH1, HNRNPK, HNRNPL, HNRNPM, KHDRBS1, KHSRP, LIN28A, LIN28B, MATR3, MBNL2, NUDT21, PABPC4, PABPN1, PCBP1, PCBP2, PTBP1, PUM1, PUM2, QKI, RBFOX1, RBFOX2, RBM47, RBPMS, SLBP, SRRM4, SRSF1, SRSF3, SRSF7, SRSF9, SRSF10, TARDBP, TIA1, TIAL1, TRA2A, YBX3, ZRANB2 |
| Non-sequence-driven | 66 | |
| RNA modification | 15 | ALKBH1, ALKBH5, FTO, METTL3, METTL14, NSUN2, PUS1, RBM15, RBM15B, WTAP, YTHDC1, YTHDC2, YTHDF1, YTHDF2, YTHDF3 |
| miRNA pathway | 10 | AGO2, DGCR8, DICER1, DROSHA, PRKRA, TARBP2, TNRC6A, TNRC6B, TNRC6C, XPO5 |
| Spliceosome | 19 | AQR, BUD13, CDC40, DDX42, EFTUD2, GEMIN5, GPKOW, PPIG, PPIL4, PRPF4, PRPF8, RBM22, SF3A3, SF3B1, SF3B4, SMNDC1, SUGP2, U2AF1, U2AF2 |
| $3'$ end machinery | 9 | CPSF1, CPSF2, CPSF3, CPSF4, CPSF7, FIP1L1, LSM11, MTPAP, WDR33 |
| Other complexes | 13 | ALYREF, CHTOP, DIS3L2, EIF4A3, EXOSC5, GTF2F1, NCBP2, NCBP3, NXF1, POLR2G, RTCB, UPF1, XRN2 |
| Fixed-target | 33 | AATF, BCCIP, DDX21, DDX51, DDX52, DDX55, DKC1, FASTKD2, FBL, GNL3, GRSF1, GRWD1, IMP3, LARP7, NIP7, NKRF, NOL12, NOLC1, NOP56, NOP58, NPM1, RPS3, RPS11, SBDS, SDAD1, SSB, SUPV3L1, TBRG4, UTP3, UTP18, WDR3, WDR43, ZNF622 |
| Mixed or unclear | 61 | ABCF1, AGGF1, AKAP1, AKAP8L, APOBEC3C, ATXN2, BCLAF1, CAPRIN1, DDX3X, DDX6, DDX24, DDX59, DHX30, EIF3A, EIF3B, EIF3D, EIF3G, EIF3H, EIF4G2, EWSR1, FAM120A, FKBP4, FMR1, FUS, FXR1, FXR2, G3BP1, HLTF, HNRNPU, HNRNPUL1, IGF2BP1, IGF2BP2, IGF2BP3, ILF3, LARP4, METAP2, MKRN1, MOV10, NIPBL, NONO, PHF6, RBM5, RBM10, SAFB, SAFB2, SERBP1, SFPQ, SLTM, SND1, STAU1, STAU2, SUB1, TAF15, UCHL5, WRN, XRCC6, YWHAG, ZC3H7B, ZC3H8, ZC3H11A, ZNF800 |

**Table S8.** Protein-macro AUROC of the six methods within groups of test examples. The three test slices are used together, and each protein is scored on the examples of its own slice. Each value is the protein-macro AUROC over the $k$ proteins that have at least 10 positive and 10 negative examples in the group. RNA length is the length of the whole spliced transcript. Bold marks the best value in each row.

| Group | $k$ | RPI-Zero | IPMiner | NPI-GNN | ZHMolGraph | CORAL | ZHMolTopoRPI |
|---|---|---|---|---|---|---|---|
| *RNA type* | | | | | | | |
| Protein-coding | 200 | **0.758** | 0.708 | 0.468 | 0.584 | 0.668 | 0.726 |
| lncRNA | 85 | **0.656** | 0.645 | 0.494 | 0.603 | 0.634 | 0.629 |
| *RNA length (nt)* | | | | | | | |
| $<$1,000 | 142 | **0.670** | 0.650 | 0.497 | 0.592 | 0.618 | 0.628 |
| 1,000–1,999 | 140 | **0.713** | 0.673 | 0.497 | 0.586 | 0.640 | 0.677 |
| 2,000–3,999 | 181 | **0.749** | 0.700 | 0.476 | 0.596 | 0.672 | 0.721 |
| 4,000–7,999 | 181 | **0.755** | 0.705 | 0.464 | 0.556 | 0.686 | 0.720 |
| $\geq$8,000 | 169 | **0.731** | 0.683 | 0.472 | 0.558 | 0.657 | 0.709 |
| *Protein class* | | | | | | | |
| Motif-driven | 48 | **0.749** | 0.686 | 0.518 | 0.616 | 0.656 | 0.715 |
| Non-sequence-driven | 65 | **0.726** | 0.698 | 0.462 | 0.574 | 0.654 | 0.694 |
| Fixed-target | 31 | **0.760** | 0.726 | 0.451 | 0.585 | 0.674 | 0.738 |
| Mixed or unclear | 60 | **0.776** | 0.721 | 0.456 | 0.584 | 0.658 | 0.736 |
| *Protein length (amino acids)* | | | | | | | |
| $<$500 | 82 | **0.743** | 0.697 | 0.458 | 0.589 | 0.662 | 0.719 |
| 500–799 | 71 | **0.758** | 0.712 | 0.478 | 0.597 | 0.682 | 0.727 |
| 800–1,250 | 35 | **0.767** | 0.716 | 0.495 | 0.568 | 0.622 | 0.713 |
| $>$1,250 | 16 | **0.730** | 0.708 | 0.464 | 0.591 | 0.620 | 0.688 |
